# PAK1 and NF2/Merlin jointly drive myelination by remodeling actin cytoskeleton in oligodendrocytes

**DOI:** 10.1101/2023.10.16.555467

**Authors:** Lucas Baudouin, Noémie Adès, Kadia Kanté, Corinne Bachelin, Hatem Hmidan, Cyrille Deboux, Radmila Panic, Rémy Ben Messaoud, Yoan Velut, Soumia Hamada, Cédric Pionneau, Kévin Duarte, Sandrine Poëa-Guyon, Jean-Vianney Barnier, Brahim Nait Oumesmar, Lamia Bouslama-Oueghlani

## Abstract

In the central nervous system (CNS), myelin formation by oligodendrocytes (OLs) relies on actin dynamics. Actin polymerization supports the ensheathment step, when the OL process contacts the axon, while a drastic shift to actin depolymerization is required to enable the following step of wrapping and expansion of myelin membranes. The molecular mechanisms triggering this switch, essential for proper myelination, have yet to be elucidated. Here, we identify P21-activated kinase 1 (PAK1) as a major regulator of actin depolymerization in OLs. We show that PAK1 accumulates in OLs in a kinase inhibited form, triggering actin disassembly and, consequently, myelin expansion. Remarkably, we identify NF2/Merlin as an endogenous inhibitor of PAK1 by proteomics analysis of its binding partners. We found that *Nf2* knockdown in OLs results in PAK1 activation and impairs myelin formation, and that pharmacological inhibition of PAK1 in *Nf2*-knockdown OLs rescues these defects. Moreover, we demonstrate that modulating PAK1 activity in OLs controls myelin expansion and provide compelling evidence indicating that specific *Pak1* loss-of-function in oligodendroglia stimulates the thickening of myelin sheaths *in vivo*. Overall, our data indicate that PAK1-NF2/Merlin duo plays a key role in actin cytoskeleton remodeling in OLs, required for proper myelin formation. These findings have broad mechanistic and therapeutic implications for demyelinating diseases and neurodevelopmental disorders.

**Significance:** Remodeling actin cytoskeleton plays a crucial role in myelin formation by oligodendrocytes (OLs). Recent studies have shown that expansion and wrapping of myelin membranes around axons depends on actin depolymerization. However, the molecular mechanisms triggering this key step in myelination are not fully elucidated. Using genetic and pharmacological tools as well as proteomics analyses, we found that PAK1 (P21 Activated Kinase 1) kinase activity is maintained inhibited by NF2/Merlin in OLs to allow actin depolymerization and, consequently, myelin membrane expansion. *Pak1* loss-of-function in OLs leads to an increase in myelin thickness in the white matter of adult mice, confirming the role of PAK1 inactivation in myelin membrane expansion.

## Introduction

Myelin is a multilayered membrane formed by oligodendrocytes (OLs) on axons in the central nervous system (CNS), increasing the conduction velocity of action potentials and providing metabolic support for neurons (1). During development, oligodendrocyte precursor cells (OPCs), emerging from the restricted ventricular regions of the CNS, migrate extensively and, ultimately, some of them differentiate into myelinating OLs (2–4). In adulthood, axons are not fully myelinated and show intermittent patterns of myelination in different regions of the CNS (5–11), allowing the formation of new myelin sheaths in response to neuronal activity or environmental cues (5, 12, 13). Myelin is also dynamic and constantly remodeled throughout life; it thickens or thins according to experience and could therefore modulate the synchronization of neuronal networks and, consequently, cognitive and motor functions (1, 14, 15).

The formation of myelin membranes and sheaths involves major morphological changes in OLs, which are supported by the remodeling of actin cytoskeleton (16–19). The first step of myelination, when the OL process extends to contact and ensheath a selected part of the axon (*i.e.,* ensheathment step), is supported by actin polymerization, while the membrane wrapping around the axon to form a multilayered myelin sheath (*i.e.,* wrapping step) is correlated with a massive actin depolymerization (16, 17). ADF/cofilin1 and gelsolin are involved in actin disassembly in OLs during the wrapping phase (16, 17). Yet the molecular mechanisms regulating the signaling pathways triggering actin depolymerization in OLs, a key process in myelination, are still unknown.

The group I of p21-activated kinases (PAKs) can control actin cytoskeleton dynamics through their kinase activity in neurons and in other cell types outside the CNS (20–25). Activation of PAK kinase activity requires a primary binding to the Rho GTPases Rac1 and Cdc42. Auto-phosphorylation of specific amino acid sites then enables full activation and efficient kinase activity (20–22, 25, 26). Phosphorylated active PAKs modulate several downstream effectors, among which they inhibit cofilin activity through its phosphorylation, thus maintaining actin in a stabilized state. Conversely, when cofilin is not phosphorylated, i.e., when PAKs are inactive, they can sever and depolymerize actin filaments (21, 27). Therefore, PAKs are reliable candidates for triggering actin depolymerization in Ols during myelin formation. Among the group I-PAKs, PAK1 and PAK3 are highly expressed in the CNS (25, 26, 28). We previously showed that PAK3 is highly expressed in OPCs but weakly expressed in OLs and that PAK3 is involved in OPC differentiation (29). Conversely, transcriptome and protein expression analyses have shown that PAK1 is highly expressed in OLs compared to OPCs (26, 30–32), suggesting that PAK1 may be a relevant candidate for controlling myelin membrane expansion, hence sheath wrapping around axons.

In the CNS, PAK1 has been mainly studied in neurons, where its involvement in several processes such as migration and dendrite/spine morphogenesis has been widely described (23, 25, 33). The function of PAK1 in CNS myelination in mammals is still unknown (26). Recently, several *de novo* activating mutations of the *Pak1* gene have been identified in children with neurodevelopmental abnormalities (intellectual disability, autism, epilepsy…). Interestingly, some of them show subcortical white matter hyperintensities on magnetic resonance imaging, which could suggest a defect in myelination (34–36). Therefore, the dysregulation of PAK1 activity may impair OL development and myelination.

In this study, we identified PAK1 as a critical modulator of actin depolymerization in OLs, controlling therefore myelin formation. We showed that PAK1 is strongly expressed in myelinating OLs, however in its inactive, unphosphorylated form. Importantly, we identified NF2/Merlin as an endogenous inhibitor of PAK1 by a proteomics analysis of its binding partners and demonstrated the functional involvement of NF2/Merlin in myelin formation. By modulating PAK1 kinase activity in OLs via pharmacological treatments or lentiviral vectors, we have shown that inhibition of PAK1 activity in mature OLs is key to the deconstruction of the actin cytoskeleton, and hence to myelin membrane formation. Interestingly, conditional deletion of *Pak1* specifically in oligodendroglia also leads to an increase in myelin thickness *in vivo*.

Overall, our study indicates that PAK1 kinase activity is inhibited in myelinating OLs, through NF2/Merlin binding, to allow proper myelin formation.

## Results

### PAK1 protein expression increases during OL maturation and is maintained inhibited

To study the role of PAK1 in myelin formation, we first assessed its expression in OLs (**Fig. 1**). We co-immunolabeled cultures of rat OLs with anti-PAK1, anti-CNPase and anti-MBP antibodies. We showed that PAK1 is strongly expressed in the soma and processes of CNP+ MBP+ mature OLs (**Fig. 1_A-B_**). In addition, STED super resolution microscopy imaging revealed PAK1 expression in the myelin membrane (**Fig. 1_B_**). We have not been able to detect PAK1 in OPCs by immunocytochemistry, while it is weakly detectable by western blot (**Sup.Fig. 1**). However, its expression is higher in OLs than in OPCs, both at the mRNA and protein levels (**Sup.Fig 1_A-B_**). We also found a consistent pattern of PAK1 expression *in vivo* in the mouse CNS white matter during development, where PAK1 is detected in Olig2+CC1+ mature OLs, but not in Olig2+CC1-OPCs (**Fig. 1_C_**), as well as in other regions of the CNS (*data not shown*). Therefore, these results show a preferential expression of PAK1 in mature OLs, arguing for its potential involvement in regulating myelin formation.

**Figure 1.**
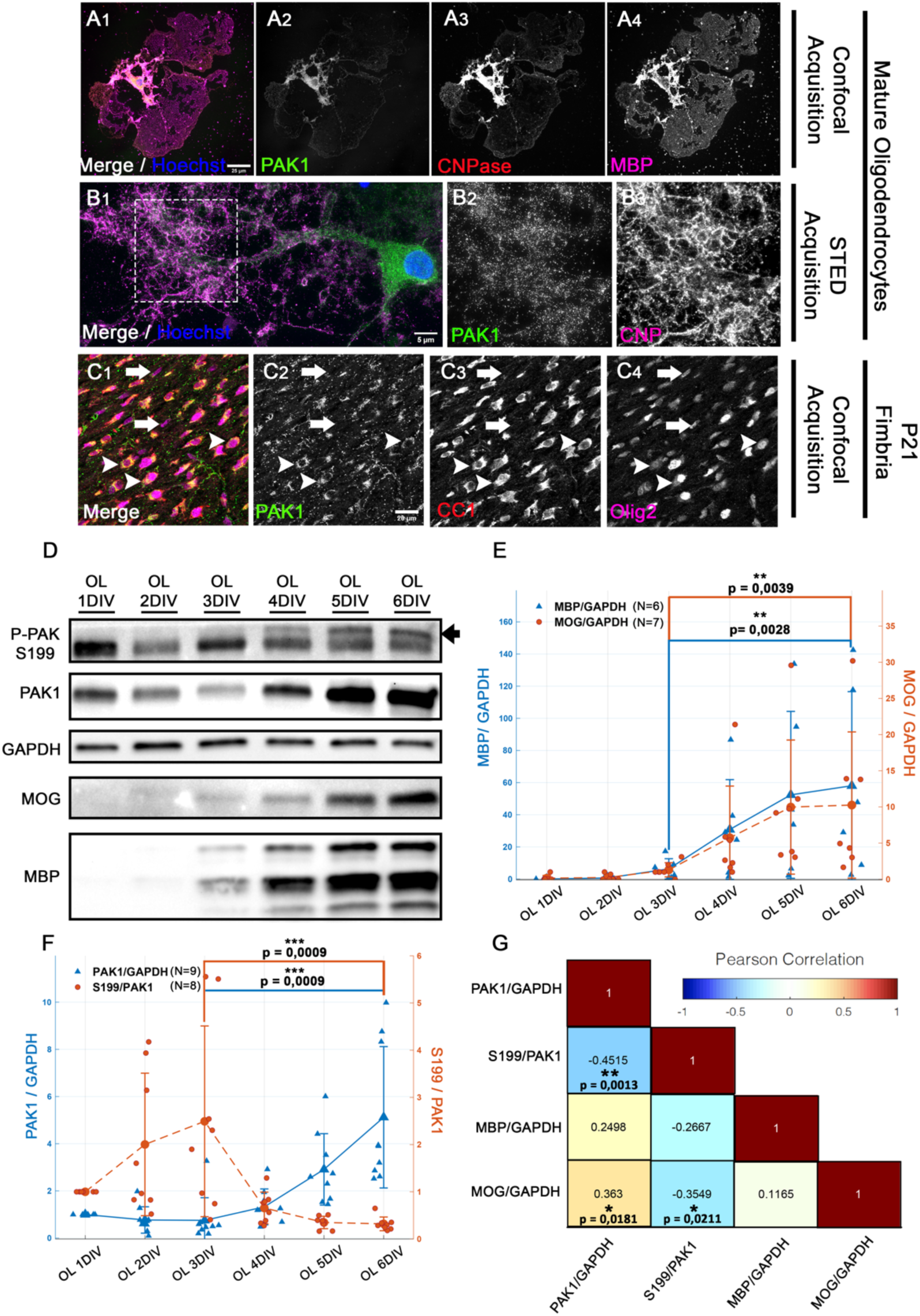
PAK1 is expressed in mature oligodendrocytes and its expression increases while its kinase activity (phosphorylation) decreases during OL maturation. (A) Confocal acquisition of a mature OL stained with Hoescht (A1) and immunostained with PAK1 (A1, A2) CNPase (A1, A3) and MBP (A1, A4) antibodies. Note that PAK1 is expressed in mature OLs with myelin membranes. Its expression is in the soma and processes of OLs. (B) STED acquisition of a mature OL stained with Hoescht (B1) and immunostained with PAK1 (B1, B2) and CNPase (B1, B3) antibodies. In the zoomed-in insert (B2, B3), note that PAK1 is present in myelin membranes. (C) Confocal acquisition of the fimbria (P21, mouse brain) immunostained with PAK1 (C1,C2), CC1 (C1, C3), and Olig2 (C1,C4) antibodies. Note that *in vivo* PAK1 expression is detected in mature OLs Olig2+CC1+ (arrowheads). However, we did not detect PAK1 expression in OPCs Olig2+/CC1-OLs (arrows). (D) Western blot illustration of P-PAK1 (PAK1 phosphorylated on serine 199, S199) and PAK1 expression in rat OL cultures at 1, 2, 3, 4, 5 and 6DIV. MBP and MOG expressions show the progressive maturation of OLs. GAPDH is used as a loading control. The arrow indicates the isoform quantified corresponding to P-PAK1. (E) Western blot quantifications of MBP (blue curve) and MOG (orange curve) in OLs from 1 to 6DIV. MBP and MOG quantities are rationalized on GAPDH and normalized on the OL 1DIV condition. Note the significant increase of both markers of mature OLs between 3DIV and 6DIV (N= number of replicates, Permutation Test, M=1000). (F) Western blot quantifications of PAK1 (blue curve) and the P-PAK1/PAK1 ratio (orange curve) in OLs from 1 to 6DIV. PAK1 and P-PAK1 quantities are rationalized on GAPDH and normalized on the 1DIV condition. Note the significant increase of PAK1 expression, while the P-PAK1 (S199)/PAK1 ratio decreases in OLs between 3DIV and 6DIV (N= number of replicates, Permutation Test, M=1000). (G) Pearson correlation matrix plots illustrating relations between the indicated normalized protein levels in OLs. Note the significant negative correlation between P-PAK1 (S199) and PAK1/MBP/MOG expression (Pearson).

As OLs in culture form flat myelin sheets whose molecular composition is similar to compact myelin *in vivo* and myelin membrane surface formed by OLs *in vitro* often mirrors myelin formation *in vivo* (16, 17, 37), we used this system to perform mechanistic analyses. Since the inhibited unphosphorylated form of PAK1 triggers actin disassembly (20, 38, 39), a process required for myelin expansion and wrapping, we hypothesized that the involvement of PAK1 in myelination could be supported by an inhibition of its kinase activity in mature OLs. To test this hypothesis, we examined PAK1 expression level and its phosphorylation state on a specific amino-acid site (serine 199, referred to as S199) in rat primary oligodendroglial cultures, from 1 to 6 days *in vitro* (DIV) in a differentiation medium (**Fig. 1_D_**). We chose to carry out this study on rat OLs, as their maturation is slower than that of the mouse, making it easier to study the different transient expressions of proteins during maturation. Phosphorylation of S199 has been shown to correlate strongly with an activated state of PAK1 and is widely used as a marker of its activity (40–45). The antibody we used also recognizes the phosphorylated forms of PAK2 and PAK3. However, their different molecular weights make them easy to recognize. Furthermore, using brain extracts from *Pak1* and *Pak3* knockout mice (data not shown), we validated our assay, and we were able to precisely quantify the phosphorylated form of PAK1. In our conditions of culture, OLs mature and form myelin membranes from 3 to 6 DIV, as assessed by the increased expression of MBP and MOG, two markers of OL maturation (16, 17, 46–48) (**Fig. 1_D,E_**). Interestingly, we found that within this time-window, PAK1 expression increases drastically, yet the proportion of its phosphorylated form decreases (**Fig. 1_D-F_**). Furthermore, we showed that the presence of active phosphorylated PAK1 is negatively correlated to PAK1 and MOG expression levels (**Fig. 1_G_**). These results indicate that OL maturation and myelin membrane formation are associated with an increased presence of PAK1 in the cell, yet in its kinase inhibited form. This large amount of inhibited PAK1 protein in mature OLs could therefore drive the depolymerization of the actin cytoskeleton required for myelin membrane expansion.

### NF2/Merlin is an endogenous inhibitor of PAK1 in OLs

The accumulation of PAK1 in an inhibited state in mature OLs strongly argues for the presence of an endogenous inhibitor of PAK1. Since PAK1 is an oncoprotein that has been widely studied in the cancer field, several inhibitors have been reported in the literature, and some of them act through binding to PAK1 (49, 50). We therefore chose to carry out a proteomics study to identify PAK1 binding partners in OLs. We immunoprecipitated PAK1 from OL cultures at 5DIV, a time within the PAK1 inhibition period (DIV3-DIV6) and used an IgG antibody as a control. We identified by mass spectrometry a total of 684 proteins in 10 independent samples (control IP and PAK1 IP, combined) (**Sup. Table1).** We also performed protein quantification and statistical analysis that allowed us to select 71 potential PAK1 binding proteins (**Sup. Table2**). The mass spectrometry proteomics data have been deposited to the ProteomeXchange Consortium via the PRIDE partner repository (51, 52) with the dataset identifier PXD044344. We performed a gene set enrichment analysis (GSEA) of all identified partners and revealed that the most abundant of them regulate cytoskeleton and intracellular signal transduction **(Sup. Fig.2)**. Out of this study, the only known PAK1 inhibitor identified was NF2/Merlin, a product of *Nf2* gene. Nevertheless, the proteomics analysis did not reveal Merlin/PAK1 interaction in all samples tested, but the interaction was significant and validated by Xtandem pipline software (**Sup. Table1**).

NF2/Merlin belongs to the ERM (Ezrin-Radixin-Moesin) protein family, and as such it transmits extracellular matrix information to the actin cytoskeleton (53–55). Both *Nf2* gene and the protein have been shown to be expressed in oligodendroglia (30, 56, 57), but the role of NF2/Merlin in CNS myelination has never been investigated. We assessed NF2/Merlin expression in mature OL cultures by co-immunolabelling of NF2/Merlin, MBP and Sox10. Our data showed NF2/Merlin expression in MBP+ OLs (**Fig. 2_A_**). Because, outside the CNS, NF2/Merlin inhibits PAK1 through a direct binding (24, 50), and in order to validate the interaction observed in the proteomics analysis, we performed immunoprecipitations of PAK1 in OLs at 4DIV (data not shown) and 5DIV (**Fig. 2_B_**) and showed that NF2/Merlin co-immunoprecipitates with PAK1 (**Fig. 2_B1_**). The reverse immunoprecipitation also showed that PAK1 co-immunoprecipitates with NF2/Merlin (**Fig. 2_B2_**). Hence, we demonstrated an endogenous interaction between PAK1 and NF2/Merlin in OLs. To determine whether this interaction results in the inhibition of PAK1 kinase activity, we compared PAK1 phosphorylation (S199) in OL cultures treated with siRNA targeting *Nf2* (si*Nf2*) or with control siRNA (siCtrl) (**Fig. 2_CD_**). si*Nf2* treatment was applied from the onset of cell differentiation, for 3 DIV, to ensure an efficient *Nf2* knockdown in OLs at 5DIV (about 60% of NF2/Merlin suppression, **Fig. 2_D1-D2_**). Remarkably, we showed that *Nf2* knockdown in OLs leads to an increased amount of active-phosphorylated S199-PAK1, without affecting the overall PAK1 expression level (**Fig. 2_E1-E3_**). Altogether, our data uncover NF2/Merlin as an endogenous inhibitor of PAK1 activity in OLs.

**Figure 2.**
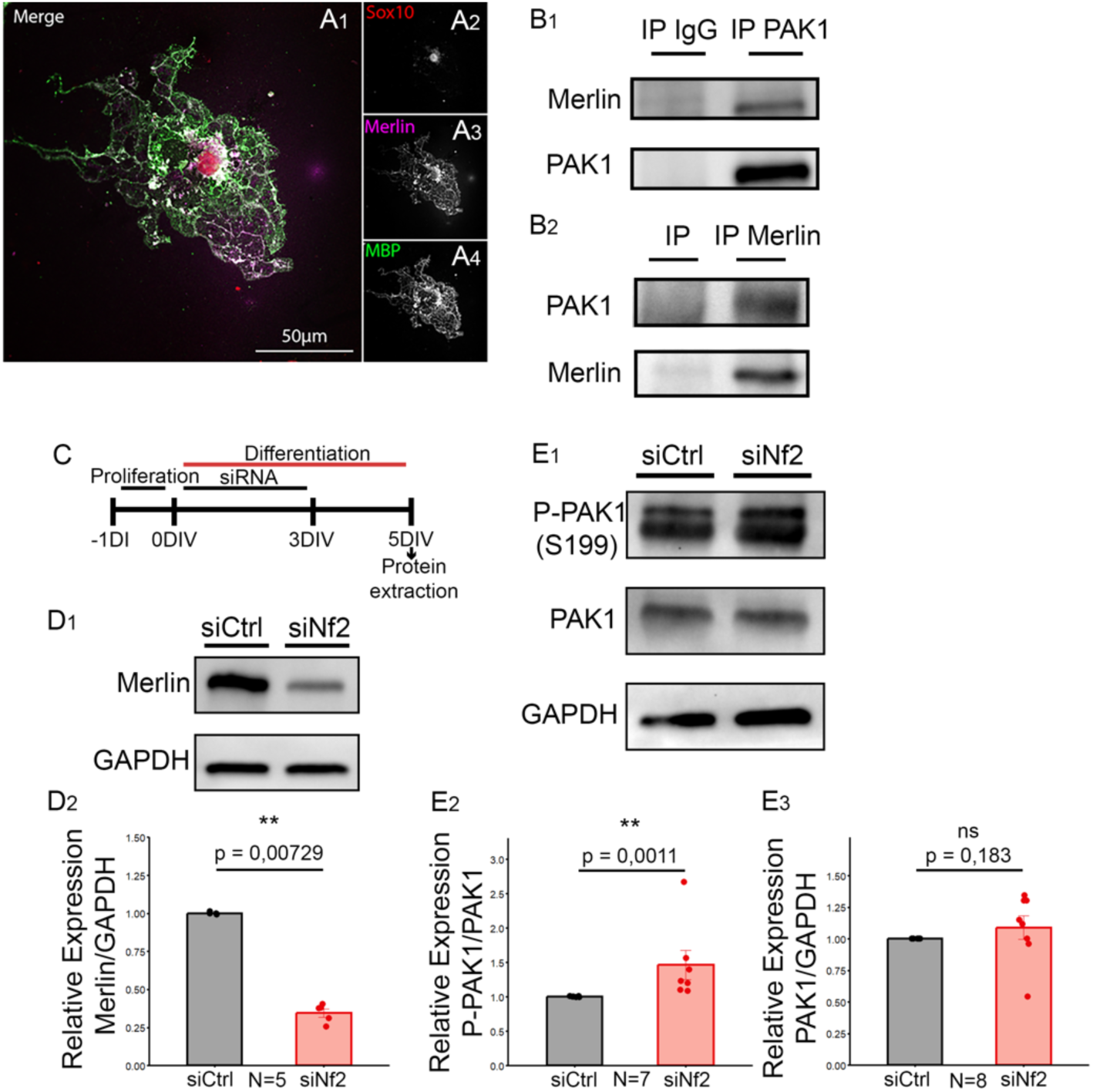
NF2/Merlin is an endogenous inhibitor of PAK1 in OLs. (A) Confocal acquisition of a mature rat OL immunostained with Sox10 (A1, A2), NF2/Merlin (A1, A3) and MBP (A1, A4) antibodies. Note that NF2/Merlin is expressed in the soma and the myelin membrane of the mature OL. (B) Western blot of PAK1 immunoprecipitation (B1) and Merlin immunoprecipitation (B2) in rat OLs at 5DIV (N=5, N=number of replicates). Note that PAK1 and NF2/Merlin co-immunoprecipitate, demonstrating the endogenous interaction between the two proteins. (C) Schematic diagram showing the different experimental steps of siRNA treatments in OL cultures. After 1 DIV in proliferation medium, siRNA treatment of OPCs is initiated at the onset of differentiation (0DIV) and left for 3DIV. After 5DIV of differentiation, cells were analyzed. (D) (D1) Western blot illustration of NF2/Merlin expression in 5DIV OLs treated with si*Nf2* or siRNA control (siCtrl). GAPDH is used as a loading control. (D2) Western blot quantification of NF2/Merlin rationalized on GAPDH and normalized on the control condition. Note that si*Nf2* treatment efficiently reduces NF2/Merlin expression in OL cultures (N= number of biological replicates, Wilcoxon rank-sum test). (E) (E1) Western blot illustration of P-PAK1 (phosphorylated on serine 199, S199) and PAK1 expression in 5DIV rat OLs treated with si*Nf2* or control siRNA. GAPDH is used as a loading control. (E2-E3) Western blot quantification of P-PAK1 rationalized on PAK1 expression (E2), and PAK1 expression rationalized on GAPDH (E3). Note that si*Nf2* treatment significantly increases PAK1 phosphorylation, and thus PAK1 catalytic activity, without affecting its expression (N= number of replicates, Wilcoxon rank-sum test).

### NF2/Merlin regulates myelin expansion through PAK1 inhibition

Having established that *Nf2* knockdown in OLs induces an increase in PAK1 activity, we next investigated whether this has any effect on myelin membrane formation. We used the surface area of myelin membranes formed by OLs as an indicator of myelin expansion. We revealed OL morphology and myelin membrane expansion by performing MBP immunostaining. We compared the myelin membrane surface formed by OLs treated with si*Nf2* and siCtrl as described in Fig. 3_A_. We found a significantly restrained myelin membrane expansion under si*Nf2* condition compared to the control at 6DIV (Fig. 3_B_). To ensure that reduced myelin membrane expansion is due to a lift of PAK1 inhibition in si*Nf2*-treated OL cultures, we performed rescue experiments using NVS-PAK1-1 (NVS), a highly specific pharmacological inhibitor of PAK1 (58, 59). We confirmed that NVS treatment effectively inhibited PAK1 kinase activity in the OL by analyzing its phosphorylation state and that of one of its downstream effectors (Fig. Sup. 3,4). Next, we confirmed that NVS treatment on si*Nf2* treated OLs did efficiently re-establish PAK1 inhibition (Sup. Fig. 4). Interestingly, we found that pharmacological inhibition of PAK1 is sufficient to rescue the defects of myelin membrane expansion observed in NF2/Merlin knockdown condition (Fig. 3_B_). To test whether NF2/Merlin-mediated inhibition of PAK1 could regulate actin cytoskeleton in OLs and subsequently myelin membrane formation, we purified F- and G-actin from OL cultures (at 6DIV), treated with si*Nf2* and siCtrl and quantified F- to F+G-actin ratios, used as an indicator of the amount of polymerized actin (16, 17). An increase in polymerized actin was observed in si*Nf2* OLs compared to controls, but NVS treatment did not restore actin state (Sup.Fig.5). However, all the treatments did not affect cell differentiation (ratio of MBP+Olig2+/Olig2+, siCtrl+DMSO 58,82%±8,24, siNf2+DMSO 63,36%±7,8, siNf2+NVS 62,75±7,63; p=0,6525, Kruskal-Wallis, N=6).

**Figure 3.**
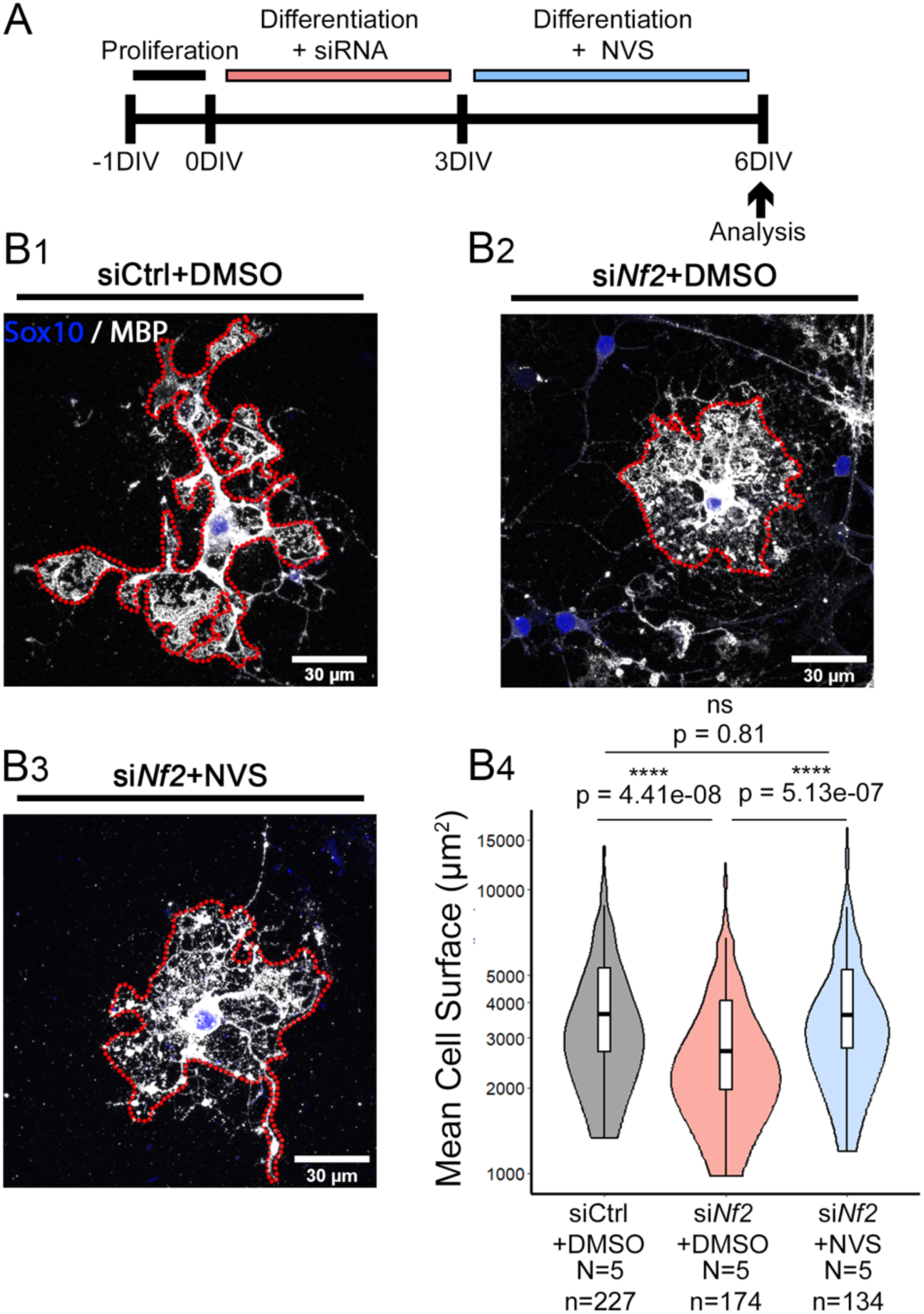
*Nf2* knockdown induces a reduction in OL myelin membrane expansion that is rescued by NVS-PAK1-1 treatment. (A) Schematic diagram showing the different experimental steps. After 1 DIV in proliferation medium, OPCs are treated with siRNA (si*Nf2* or control siRNA (siCtrl)) for 3 DIV in differentiation medium. At 3DIV, medium is changed and DMSO or NVS (250nM) is added. After 3DIV of NVS or DMSO treatment (6DIV of differentiation), cells were analyzed. (B) Confocal acquisition of OLs treated with siCtrl and DMSO (B1), si*Nf2* and DMSO (B2) or si*Nf2* and NVS (B3) and immunostained with MBP (white) and Sox10 (blue) antibodies. The red dotted lines delimit the MBP+ surface used for quantifying the myelin membrane surface. (B4) Quantification of the MBP+ cell surface in OLs treated with siCtrl and DMSO (grey), si*Nf2* and DMSO (red), or si*Nf2* and NVS (blue). Note that Si*Nf2* and DMSO treatment reduces myelin membrane surface compared to the control condition (siCtrl and DMSO). Note that re-establishing PAK1 inhibition with NVS treatment in the si*Nf2* condition normalizes myelin membrane expansion (N= number of replicates, n= number of cells, Kruskal-Wallis followed by Dunn post-hoc test).

Altogether, these findings demonstrate that NF2/Merlin inhibition of PAK1 activity in OLs plays a critical role in myelin membrane expansion.

### Constitutively active PAK1 restrains myelin membrane expansion through actin cytoskeleton assembly

To further confirm the need for PAK1 inhibition to induce myelin membrane expansion, it is important to directly manipulate PAK1 activity and determine its effects on OL membranes and the actin cytoskeleton. To do so, we specifically modulated PAK1 activation state in OL cultures. First, we analyzed the effects of constitutively active PAK1 in counteracting endogenous PAK1 inhibition. As specific pharmacological activators of PAK1 are not available to date, we used inducible lentiviral vectors to overexpress a constitutively activated form of PAK1 (PAK1 CA, T423E) in oligodendroglia. We generated doxycycline-inducible lentiviral vectors that express a Kusabira-Orange, fluorescent reporter, to assess cell transduction efficiency and a HA-tagged PAK1 CA or GFP (used as a control) (**Fig. 4_A_**). We validated these vectors in HEK-293T cell line, as the rate of lentiviral transduction in these cells is much higher than that of OLs and are therefore more suitable for western blot analysis. After assessing the increased amount of phosphorylated PAK1 in PAK1 CA cultures (**Sup. Fig. 6**), we analyzed whether PAK1 CA was efficient for modulating actin cytoskeleton. To do so, we purified F- and G-actin fractions from cells over-expressing PAK1 CA or GFP, and quantified F- to F+G-actin ratios. As expected, PAK1 CA-transduced cells showed a significant increase of the fraction of F-actin compared to control (**Fig. 4_B_**), indicating the maintenance of actin in a preferential polymerized state. Next, we assessed the effects of these vectors on primary oligodendroglial cultures as described in **Fig. 4_C_**. We analyzed only transduced (Kusabira+) and induced (GFP+ or HA+) mature MBP+ OLs (MBP+) (**Fig. 4_D_**). Interestingly and consistent with the maintenance of actin in an abnormal polymerized state, we found that the myelin membrane surface was significantly reduced in PAK1 CA-transduced OLs compared to controls at 4DIV (**Fig. 4_E_**). The percentage of differentiated OLs in these cultures was not affected in PAK1 CA condition compared to the control cultures (39%±8 versus 41%±8, p=0.841, Wilcoxon - Mann Whitney, N= 5). Overall, these results indicate that maintenance of PAK1 in an abnormal active state in mature OLs specifically disturbs myelin membrane expansion, through the regulation of actin cytoskeleton.

**Figure 4.**
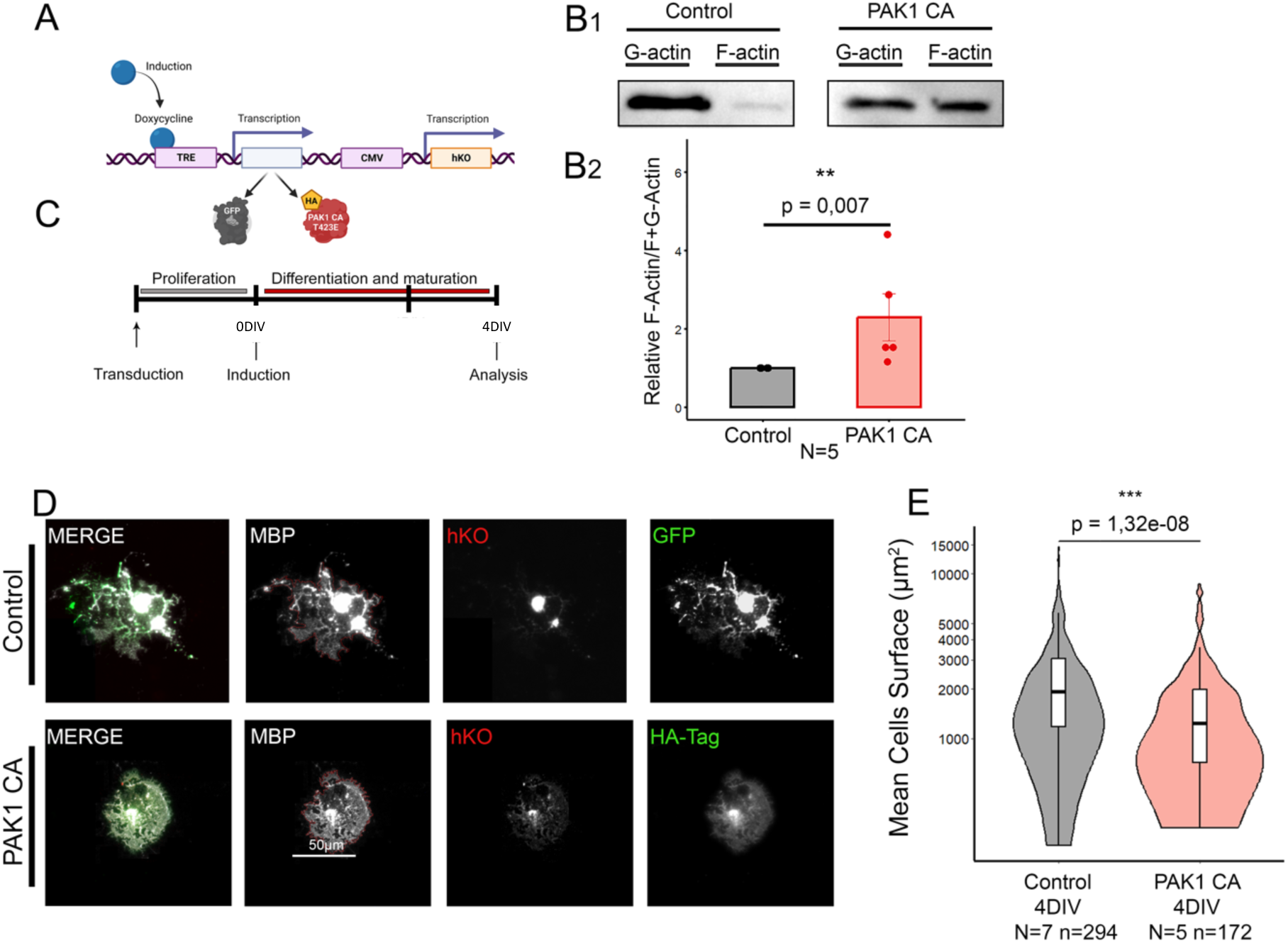
Constitutively active PAK1 maintains actin polymerized and restrains myelin membrane formation. (A) Schematic representation of the lentiviral vectors used in this study: doxycycline inducible lentiviral vectors expressing GFP (control) and a mutant form of PAK1 (HA-tagged) leading to its constitutive activation (PAK1 CA, T423E). (B) (B1) Western blot analysis of purified F- and G-actin from HEK-293T cells transduced with control or PAK1 CA lentiviruses. (B2) Quantification of the ratio of F-actin over total actin (F+G-actin) in PAK1 CA and control conditions. Note the significant increase of F-actin/total actin ratio under PAK1 CA condition compared to control, indicating that PAK1 CA induction favors the maintenance of actin in a polymerized state (N= number of biological replicates, Wilcoxon rank-sum test). (C) Schematic diagram of the transduction protocol in primary rat OPC cultures. Viral transduction is performed in proliferating OPC cultures. Inductions of GFP and PAK1 CA are induced at the onset of differentiation (0DIV), cells are then analyzed at 4DIV. (D) Mature OLs transduced with control or PAK1 CA vectors are assessed by the expression of the Kusabira-Orange (hKO) (red). Induction of viral expression is assessed by the GFP or HA (green) expression. MBP staining (white) allows the quantification of the myelin membrane surface (red dotted line). (E) Cell surface quantification of OLs transduced with control or PAK1 CA vectors and analyzed at 4 DIV. Note that PAK1 CA induction significantly limits membrane surface expansion compared to control (N= number of replicates, n= number of cells, Wilcoxon rank-sum test).

### PAK1 inhibition increases myelin membrane expansion and myelination *ex vivo* via actin disassembly

We then tested whether specific reinforcement of PAK1 inhibition in OL cultures would affect the actin cytoskeleton and, consequently, membrane expansion. To this aim, we treated oligodendroglial cultures with NVS during the first three days of differentiation, when PAK1 expression starts to increase and prior to the onset of its endogenous inhibition (**Fig. 5_A_**). To assess the effects of NVS-mediated PAK1 inhibition on actin cytoskeleton, we purified F- and G-actin fractions from OLs treated with NVS and DMSO (for control condition), as described above, and performed analysis at 3DIV and 6DIV. In line with our hypothesis, we found a greater decrease of polymerized actin under PAK1 inhibition at 3 DIV compared to control (**Fig. 5_B_**), indicating that PAK1 inhibition favors actin depolymerization in early maturing OLs. However, in fully mature OLs, even though we observed a tendency to a decreased polymerized actin at 6 DIV, it did not reach significance (1±0 versus 0,852±0,20, p= 0.0820, Wilcoxon - Mann Whitney, N=8). This could be explained by the almost completely depolymerized actin in the control condition at 6 DIV, therefore reducing the difference with the NVS condition (17). To determine if the increase of actin depolymerization observed is correlated to an increase of myelin membrane expansion, we performed immunostaining for MBP in OL cultures at 3 and 6 DIV under DMSO or NVS treatments (**Fig. 5_C-D_**). We found that enhancing PAK1 inhibition with NVS treatment significantly increased myelin membrane expansion both at 3 (**Fig. 5_E1_**) and 6 DIV (**Fig. 5_E2_**). The percentage of differentiated OLs in these cultures was not affected by NVS treatments (*at 3DIV: 38%*±2 versus 39%±8, p=0,69, at 6DIV: 57%±5 versus 57%±4, p=0,69, Wilcoxon - Mann Whitney, N=5). In line with these results, we showed that overexpressing a kinase dead form of PAK1 (inducible lentiviral vector PAK1 KD, K299R) in OLs leads to an increase of membrane surface at 4DIV (**Sup.Fig. 6_B_**) without affecting cell differentiation (39%±8 versus 38%±12, p=0.904, Wilcoxon - Mann Whitney, N=5). Overall, these results show that PAK1 inhibition in OLs stimulates specifically membrane expansion by enhancing actin depolymerization.

**Figure 5.**
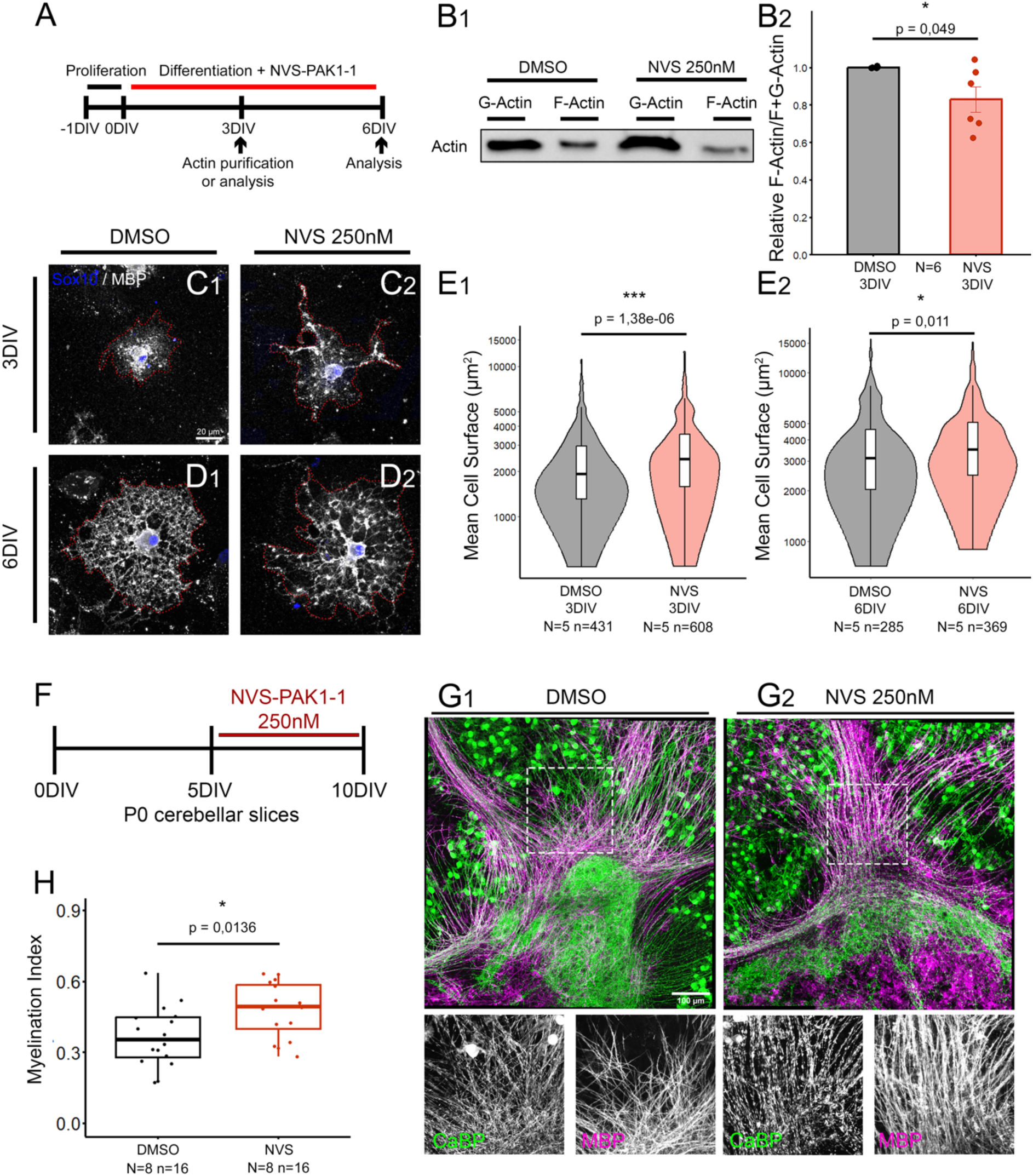
Pharmacological inhibition of PAK1 (NVS-PAK1-1) enhances actin depolymerization and consequently myelin membrane expansion and myelination *ex vivo*. (A) Schematic diagram showing the different experimental steps of NVS-PAK1-1 (NVS) treatment on primary rat OPC cultures. After one day in a proliferating medium, NVS treatment is initiated at the onset of differentiation. Differentiated OLs are then analyzed at 3 and 6 DIV. DMSO is used as a vehicle control. (B) (B1) Western blot analysis of F- and G-actin fractions purified from OLs treated with DMSO or NVS 250nM at 3 DIV. (B2) Quantification of the ratio of F-actin over total actin (F+G-actin) in OLs at 3DIV. Note the significant decrease of F-actin /total actin ratio in the NVS condition compared to control (N= number of replicates, Wilcoxon rank-sum test). (C-D) Mature OLs treated with DMSO (C1, D1) or NVS 250nM (C2, D2), stained with MBP (white) and Sox10 (blue) and analyzed at 3DIV (C) and 6DIV (D). The MBP+ surface of OLs is delimited by the red dotted line, which is used to quantify myelin membrane surface area. (E) MBP+ surface quantification of OLs treated with DMSO or NVS 250nM and analyzed at 3 (E1) and 6 DIV (E2). Note the significant increase of myelin membrane surface in the NVS condition compared to control at 3 and 6DIV (N= number of replicates, n= number of cells, Wilcoxon rank-sum test). (F) Schematic diagram showing the different experimental steps of NVS-PAK1-1 (NVS) treatment on organotypic cultures of P0 mouse cerebellar slices. After 5 DIV, NVS treatment is initiated on cerebellar slices for 5 DIV (from 5 to 10 DIV). DMSO is used as a control vehicle. (G) Confocal acquisition of CaBP (green, Purkinje cells) and MBP (magenta, OL and myelin) immunostaining illustrating myelination of cerebellar cultures treated with DMSO (G1) or NVS 250nM (G2). The two inserts on the bottom of each image represent a zoom in of CaBP and MBP staining. (H) Boxplot of the quantification of myelination by calculating the index of myelination. Note the significant increase of this index in the NVS condition compared to DMSO treated slices, indicating that strengthening PAK1 inhibition increases myelination (N= number of replicates, n= number of cerebellar slices, unpaired Student’s t-test).

Next, to determine whether PAK1 inhibition could regulate axonal myelination, we used organotypic cerebellar slices (60). We treated newborn cerebellar slices with NVS or DMSO from 5 to 10DIV (**Fig. 5_F_**), a period of intense OL differentiation and myelination (60), and stained Purkinje cells and myelin with CaBP and MBP antibodies, respectively (**Fig. 5_G_**). We then quantified myelinated axons and calculated the myelination index, as previously described (60). In line with our *in vitro* data, sustained PAK1 inhibition with NVS treatment in organotypic cerebellar slices significantly enhanced myelination (**Fig. 5_G-H_**). As the increase of myelination could be related to a potential enhancement of oligodendroglial cell number, which in turn will form more myelin internodes, we quantified their differentiation and density in slices immunostained for Sox10 and CC1 (**Fig.Sup. 7**). The density of differentiated OLs was not significantly different between control and NVS treated slices (**Fig.Sup. 7**). Even though we could not exclude that the observed effects of NVS could be related to PAK1 inhibition in neurons and other glial cells (25, 26), our results indicate that sustained inhibition of PAK1 activity in OLs promotes myelination *ex vivo*.

### *Pak1* deletion in OLs induces an increase of myelin thickness *in vivo*

To further extend our findings to *in vivo*, we generated a conditional knockout mouse with *Pak1* deletion specifically in oligodendroglia by crossing Cnp-Cre and *Pak1*-floxed mice. These animals were also crossed with Rosa26-YFP mice to identify recombined cells. We therefore generated in this study *Pak1^flox/flox^;*CNP^cre/+^*;Rosa-YFP* mice, hereafter referred to as *Pak1*cKO and *Pak1^flox/flox^;*CNP^+/+^*;Rosa-YFP* mice as control animals. We assessed the rate of Cnp promoter driven Cre recombination in white matter and confirmed that up to 80% of Olig2+ cells are YFP+ (*data not shown*), as described previously (17, 61). We also validated the decreased PAK1 expression in the optic nerve of *Pak1*cKO mice compared to controls (**Fig. 6_A_**). The remaining PAK1 protein detected in *Pak1*cKO optic nerves is probably due to its expression in other cell types, including neurons. We then showed that the percentage of differentiated OLs (Olig2+/CC1+ OLs among the total Olig2+) remained equally unchanged between controls and *Pak1*cKO (**Fig. 6_B_**). Next, we examined the effect of *Pak1* deletion on myelin integrity and thickness *in vivo* by electron microscopy in adult white matter regions at P60 (**Fig. 6_C1-C2_**). In the anterior commissure of *Pak1*cKO, most of axons presented thicker myelin, as assessed by the significantly decreased g-ratio (with respect to control (**Fig. 6_D_**). Scatter plots of the g-ratio as a function of axon diameter showed that myelin thickness was higher across the whole spectrum of axon diameters in the *Pak1*cKO (**Fig. 6_E_**), indicating a global myelin thickening rather than an axonal diameter-dependent phenotype. In addition, we did not observe significant changes in the density of myelinated axons (**Fig. 6_F_**) nor in axonal diameters (**Fig. 6_G_**). We obtained similar results in the corpus callosum with a decrease in the g-ratio in *Pak1*cKO mice and no change in myelinated axon density or axon diameter (**Sup. Fig 8**). Interestingly, OL cultures lacking PAK1 expression had larger membrane areas than control OLs (**Sup. Fig 9**), while retaining the same rate of differentiation (18%±5 versus 11±2, p= 0.309, Wilcoxon - Mann Whitney, N=5). These results confirm that myelin membrane expansion *in vitro* recapitulates myelin thickening *in vivo*.

**Figure 6.**
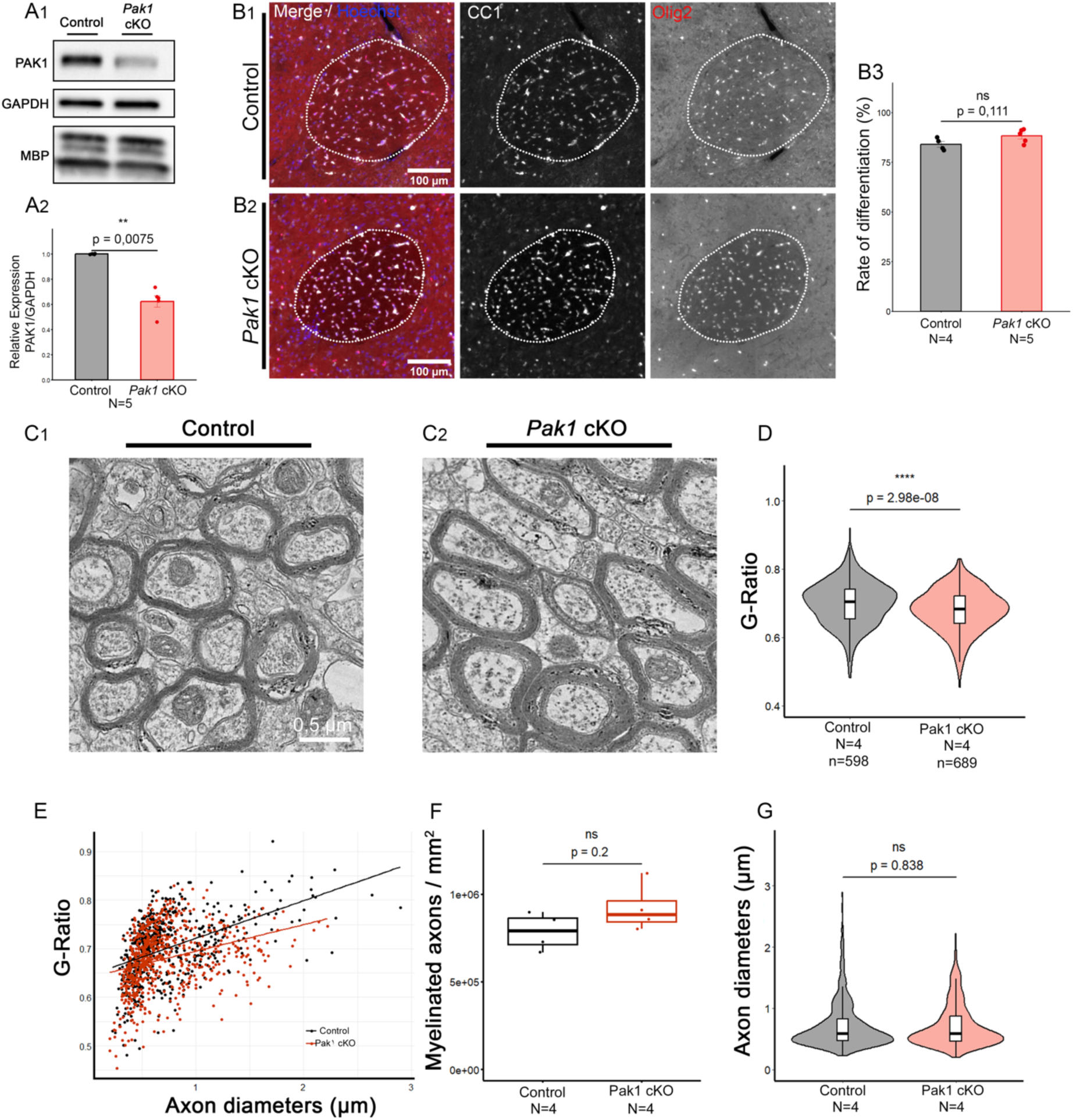
Myelin thickness is increased in the white matter of *Pak1* cKO adult brain. (A1) Western blot illustration of PAK1 and MBP expression in control and *Pak1* cKO optic nerve lysates at P22. GAPDH is used as a loading control. (A2) Western blot quantification of PAK1 expression rationalized on GAPDH and normalized on the control condition. Note the significant decrease of PAK1 expression in *Pak1* cKO optic nerve compared to control (N= number of replicates, Wilcoxon rank-sum test). (B) Axioscan imaging of the anterior commissure at P60 in control (B1) and *Pak1*cKO (B2) animals. The anterior commissure is indicated by the dotted white line. Differentiated OLs (CC1+) are shown in white and total oligodendroglial cells (Olig2+) in red. (B3) Quantification of the percentage of differentiated OLs (CC1+Olig2+ cells over the Olig2+ cell population) in the anterior commissure at P60 in control and *Pak1*cKO animals. Note that the percentage of differentiated OLs is not different between the two genotypes (Mann Whitney test). (C) Electron micrographs of transverse myelinated axons in the anterior commissure of young adults (P60) control (C1) and *Pak1*cKO mice (C2). (D) Boxplot of g-ratio analysis in control and *Pak1*cKO mice. Note the significant decrease in the g-ratio in *Pak1*cKO animals compared to controls, indicating an increase in myelin thickness in the *Pak1*cKO anterior commissure (N= number of animals, n= number of myelinated axons quantified, unpaired Student’s t-test). (E) Scatterplots of individual axon diameter measurements as a function of axon g-ratio (each individual data point represents one axon) and linkage to the regression line for each genotype. (F) Boxplot of myelinated axon densities. Note that *Pak1* deletion in OLs did not affect the density of myelinated axons (N= number of animals, Wilcoxon rank-sum test). (G) Boxplot of axon diameters. Note that *Pak1* deletion in OLs did not affect axonal diameter of neurons (N= number of animals, Wilcoxon rank-sum test).

Taken together, these results indicate that loss-of-function of *Pak1 in vivo* in oligodendroglia promotes myelin thickening in white matter, further supporting the *in vitro* data demonstrating that PAK1 deletion/inhibition stimulates myelin membrane expansion.

## Discussion

Understanding the molecular mechanisms regulating myelin formation is essential, given the adaptive properties of this structure and its crucial roles in CNS function. We used complementary genetic and pharmacological tools to show that proper myelin formation relies on the endogenous inhibition of PAK1 activity in OLs, which in turn triggers actin disassembly. Importantly, we identified NF2/Merlin as an endogenous inhibitor of PAK1 in OLs, and this inhibition is essential for actin disassembly and subsequent myelin formation (**Fig. 7**).

**Figure 7.**
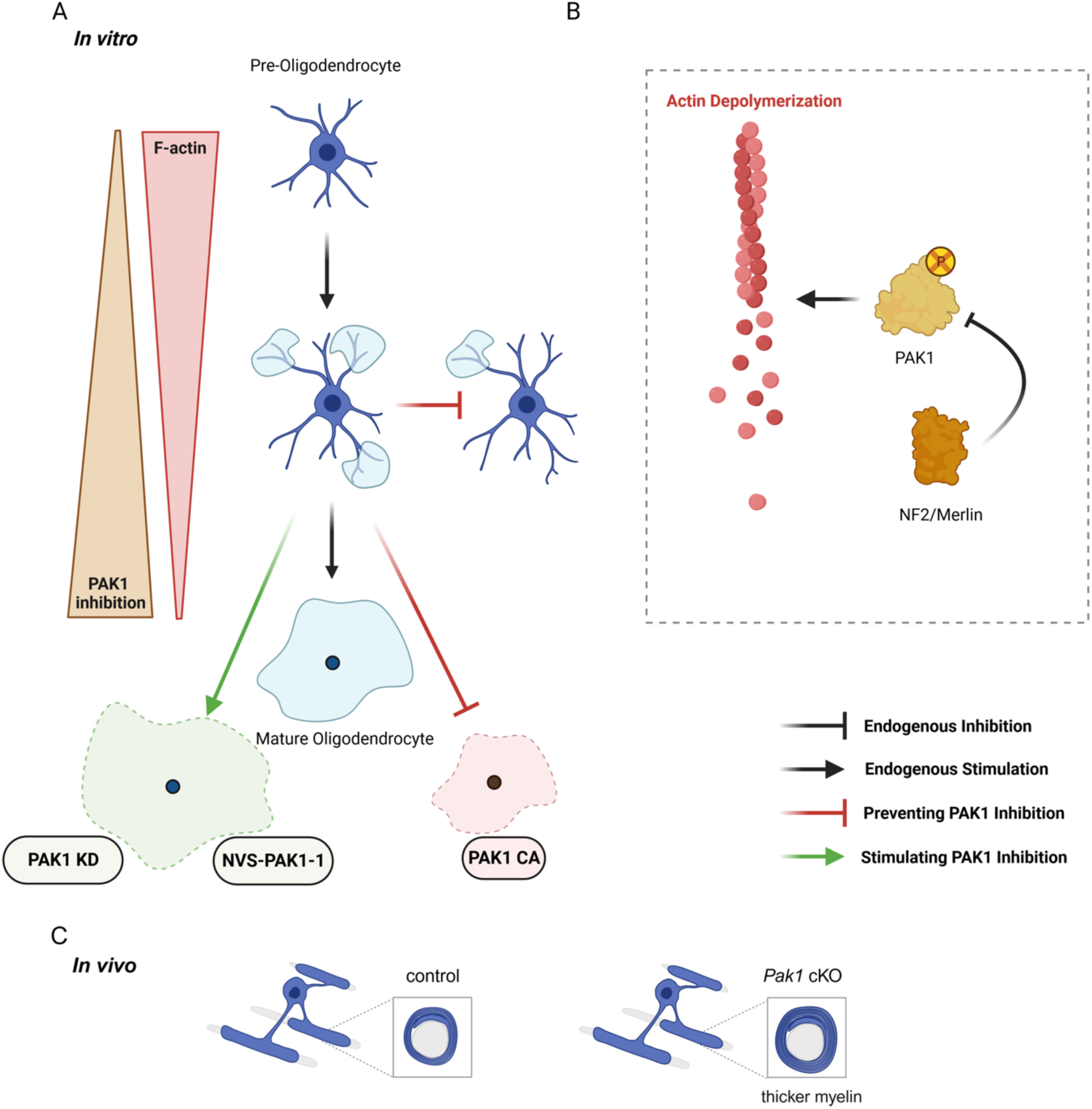
Diagram showing the proposed model of actin cytoskeleton regulation by the PAK1-NF2/Merlin duo during myelin membrane formation. (A) During *in vitro* OL maturation, as actin filaments depolymerize, PAK1 levels increase in tandem with its inhibition. Modulation of PAK1 kinase activity regulates the expansion of OL membranes: prevention of endogenous PAK1 inhibition limits myelin membrane formation and expansion (red inhibition arrows, PAK1 CA experiments); whereas strengthening its inhibition stimulates and increases the expansion of myelin membranes (green stimulation arrows, PAK1 KD and NVS experiments). (B) PAK1 inhibition in OLs allows actin depolymerization. NF2/Merlin is identified as an endogenous inhibitor of PAK1 in OLs. (C) Constitutive and conditional deletion of *Pak1* in OLs (*Pak1*cKO) leads to an increase in myelin sheath thickness *in vivo*.

The role of PAK1 and NF2/Merlin in mammalian myelination and their functional interaction in OLs have not been previously documented. A recent study provided evidence indicating a positive role for PAK1 in myelin formation in the zebrafish (32), unlike what we have observed in rodents. This discrepancy could be related to the fact that group I PAKs are different in zebrafish. Indeed, not only do zebrafish lack the *Pak3* gene, but it has also been shown that *Pak1* mRNA accumulates in OPCs and not in OLs in the zebrafish (62), which is the opposite in rodents, as described in our study. It is possible that some of the pro-differentiation functions associated with the mammalian PAK3 (29), which may affect the myelination process, are assigned to the zebrafish PAK1. In the same study (32),the authors also investigated the effect of group I PAK inhibition (FRAX486) on the actin cytoskeleton of mouse OPCs, but not on OLs.

We have shown that PAK1 expression increases in an inhibited unphosphorylated state, triggering the actin disassembly required for myelin membrane expansion. It is intriguing that a protein kinase is produced in large quantities to remain inhibited. However, large amounts of PAK1 proteins are possibly required for a specific and faster regulation of actin cytoskeleton dynamics to ensure rapid myelin thinning during the process of adaptive myelination. In addition, PAK1 plays other cellular roles as a scaffolding protein, i.e., independently of its kinase activity, as has been described in other cell types (63, 64). PAK1 would therefore have other roles in OLs that it would be interesting to identify in the future. In line with this assumption, our proteomics analysis revealed several potential PAK1 binding partners that are known to control a wide range of cellular functions such as protein translation and intracellular membrane trafficking (**Sup. Table 2**).

Interestingly, we found that NF2/Merlin is an endogenous inhibitor of PAK1 in OLs and is therefore required for myelin expansion. Nevertheless, NF2/Merlin plays multiple cellular roles, including direct regulation of the actin cytoskeleton independently of PAK1. Indeed, NF2/Merlin is known to stabilize actin filaments and its deletion leads to actin depolymerization in other cell types (65–68). It is therefore possible that the restricted myelin membrane formation observed upon *Nf2* knockdown is the result of multiple altered pathways. Interestingly, restoration of PAK1 inhibition with NVS treatment fully restored the ability of OLs to increase myelin membrane formation, supporting the hypothesis that NF2/Merlin mediates myelination at least partially through its action on PAK1. However, we cannot exclude another possible outcome of this result, namely that PAK1 inhibition is an extremely powerful lever controlling myelination. In this case, even if NF2/Merlin regulation of myelination was only partially mediated by PAK1, inhibiting PAK1 would restore myelination. However, surprisingly, NVS treatment did not restore polymerized actin in *siNf2* OLs, although this may be due to the sensitivity of the technique. It will therefore be important in future studies to decipher the role of NF2/Merlin in myelination *in vivo*, as well as its other mechanisms of action in OLs.

Specifically preventing PAK1 inhibition by expressing a constitutively active PAK1 mutant results in a smaller surface area of myelin membranes. Thus, PAK1 activation in OLs inhibits myelin membrane expansion. In addition, enhancing PAK1 inhibition (NVS treatment or PAK1 KD expression) as soon as PAK1 expression occurs, leads to an increase in myelin membrane surface area. Therefore, PAK1 is a major regulator of actin depolymerization in OLs and, consequently, of myelin membrane expansion. However, other factors are involved in the completion of the steps relying on actin polymerization, *i.e.,* process formation and ensheathment. Indeed, WAVE1 and N-WASP, two proteins known to activate actin polymerization via the Arp2/3 complex, have been shown to be involved in these processes (69, 70). However, the course of their activation/inactivation during OL differentiation/maturation remains unknown, like that of their upstream regulators, the Rho GTPases Rac1 and Cdc42.

Many downstream effectors of PAK1 leading to actin disassembly are known, including cofilin (20, 38). Consistently, we detected a decrease of cofilin phosphorylation in OLs treated with NVS (**Sup. Fig. 5**). Moreover, proteomics analysis of PAK1 potential partners revealed that cofilin-1 was one of them **(Sup. Table 2)**, confirming the involvement of these two proteins in a common OL signaling pathway.

Interestingly, deletion of PAK1 expression in OLs results in increased myelin membrane expansion, as in OLs with inhibited activity. This result suggests that PAK1 is required to adjust myelin membrane expansion and confirms that PAK1 inhibition promotes myelin formation. In line with this result, specific deletion of *Pak1* in oligodendroglia *in vivo* leads to an increase in myelin thickness. It would be interesting to test the effect of pharmacological inhibition of PAK1 *in vivo*, particularly under conditions of demyelination. The use of NVS-PAK1-1 is very complex *in vivo* due to the short half-life of the molecule and the need to add pharmacokinetic inhibitors to prevent degradation (59). However, a new version of this molecule has recently been generated, a PAK1 selective degrader, and has been validated *in vitro* (71). This compound could therefore be tested in future *in vivo* studies.

Importantly, mutations in the Pak1 gene, which induce kinase activation of the protein, lead to developmental disorders such as ASD, and some are associated with white matter alterations (34–36). It would therefore be interesting to determine, in animal models of PAK1 activation, the consequences on CNS myelination.

In conclusion, we have highlighted a new mechanism for myelination via regulation of the PAK1-NF2/Merlin duo. We propose that PAK1 inhibition by NF2/Merlin in OLs leads to actin depolymerization and consequently to myelin membrane expansion (**Fig. 7**). Importantly, our study opens new avenues for the understanding of the mechanisms of CNS myelination, it also provides new opportunities for translational research in myelin diseases such as multiple sclerosis, as well as in neurodevelopmental disorders where PAK1 activity is impaired.

## MATERIAL AND METHODS

### Animals

Swiss mice and Wistar rats (Janvier Labs, Le Genest-Saint-Isle, France) were used in this study. The experimental plan was designed in accordance with the European Union Guidelines for the care and use of experimental animals. They were approved by the French ethical committee for animal care of the ICM and the Ministry of National Education and Research (Project N° 2017092519011284). Mutant mice were bred and maintained on a C57BL/6N background. *Cnp*-Cre mice (61) and *Rosa26*-EYFP (72) mice have been described previously. *Pak1^Flox^* (C57BL/6N-Pak1^tm1c(EUCOMM)Hmgu^/H) mice were obtained from the MRC Harwell Institute which distributes these mice on behalf of the European Mouse Mutant Archive (www.infrafrontier.eu<u>; Repository number EM:10463</u>) with 2 LoxP sites flanking exon 5 of *Pak1*. Male *Pak1* cKO (Pak1^flox/flox^ ;Cnp ^Cre/+^ ;Rosa-YFP ^+/-^) and control animals (Pak1 ^flox/flox^ ;Cnp ^+/+^;Rosa-YFP ^+/-^) were used in this study.

### Primary rodent OPC culture

Primary cultures of glial cells were obtained from forebrain animals at P1–P2, following a protocol described in (60). Tissue was dissociated mechanically until homogenization in DMEM GlutaMAX (Invitrogen, Villebon-sur-Yvette, France) containing 10% of fetal bovine serum (Invitrogen), 100 U/mL penicillin–streptomycin (Life Technologies, Waltham, USA). Cells were plated on polyornithine (10 μg/mL, Sigma-Aldrich, St. Quentin Fallavier, France) coated flasks. Cultures were maintained at 37°C with 5% CO2 and the medium was changed the fifth day of culture and then every 2 days. OPC cultures were obtained from primary glial cultures of 8–10DIV. The first step of shaking was performed (1 hr, 250 rotations per min, 37°C) to eliminate microglial cells. To detach OPCs from the astrocyte layer, a second longer shaking step was performed (18 hr, 250 rotations per min, 37°C). The supernatant is then recovered and several steps of preferential adhesions (4 × 10 min on Falcon Petri dishes, 37°C, 5% CO2) were performed to remove remaining microglia and astrocytes. Last, a 5 min centrifugation at 1000 rpm segregated OPCs and allowed a resuspension of cells in fresh DMEM GlutaMAX, 10% of fetal bovine serum, 100 U/mL penicillin–streptomycin medium. Finally, OPCs were counted and plated in polyornithine (20 μg/mL, Sigma-Aldrich, St. Quentin Fallavier, France) coated dishes or coated 12mm glass coverslips.

### Viral transduction

Lentiviral vectors pCSIV-eGFP-IRES-hKO (control), pCSIV-PAK1-K299R-IRES-hKO (PAK1 mutant kinase dead - KD) and pCSIV-PAK1-T423E-IRES-hKO (PAK1 mutant constitutively active - CA) were used in this study. These vectors allowed doxycycline inducible expression of enhanced green fluorescent protein (eGFP) (60) and PAK1 mutants, through a rtTA (reverse tetracycline controlled transactivator)-dependent system. Transduced cells express the human Kusabira-Orange (hKO) fluorescent protein under the CMV (60). These vectors were constructed by inserting the eGFP/PAK1 mutant sequence by recombination, using the Gateway system into the pCSIV-IRES-hKO plasmid provided by Dr Hyroyushi Miyoshi (Keio University). OPCs were transduced at an MOI of 10, supplemented with 0.5µg/ml Polybren (Sigma-Aldrich, St. Quentin Fallavier, France) to increase the efficiency of gene transfer. OPCs are kept in a proliferating medium (DMEM GlutaMAX (Life Technologies)/2% B27 (Life Technologies)/1% antibiotics (Penicillin–Streptomycin [10,000 U/ mL], Sigma-Aldrich)/bFGF (25 ng/mL, Sigma-Aldrich, St. Quentin Fallavier, France)) and PDGFbb (10 ng/mL, Sigma-Aldrich, St. Quentin Fallavier, France)). eGFP and PAK1 mutant expression were induced with doxycycline (2 μg/mL, Sigma-Aldrich) at 1DIV post-transduction. At 2DIV post-transduction, cells were cultured in differentiation medium (DMEM GlutaMAX (Life Technologies)/2% B27 (Life Technologies)/1% antibiotics (Penicillin–Streptomycin [10,000 U/ mL], Sigma-Aldrich, St. Quentin Fallavier, France)/ 40 ng/ml of T3 thyroid hormone (Sigma-Aldrich, St. Quentin Fallavier, France)).

HEK 293-T cells were also transduced to validate the vectors and their effects on PAK1 expression and activation level. HEK 293-T cells were seeded in 100 mm Petri dishes (7.5 10^5^ cells/ dish) and transduced with the lentiviruses (MOI 5) supplemented with Polybren (0.5 μg/mL, Sigma-Aldrich). Transduced HEK cells were kept in a proliferating medium (DMEM GlutaMAX (Life Technologies), 10% of foetal bovine serum (Invitrogen, Villebon-sur-Yvette, France), 1% antibiotics (Penicillin– Streptomycin [10,000 U/ mL], Sigma-Aldrich, St. Quentin Fallavier, France), 1% MENAA (Invitrogen, Villebon-sur-Yvette, France)). Expression of the eGFP protein and the PAK1 mutant was induced by doxycycline (2 μg/mL, Sigma-Aldrich, St. Quentin Fallavier, France, St. Quentin Fallavier, France). Cells were lysed 3DIV after viral induction before they reached confluence.

### Quantitative Real time reverse transcription-PCR

Cultured cells were lysed at 4°C in RLT buffer (Qiagen, Courtabœuf, France) and stored at -80°C. RNAs were extracted from frozen samples using RNeasy Plus Mini kit (Qiagen, Courtabœuf, France) as described by the manufacturer. RNA concentrations were quantified with a nanodrop (Thermoscientific). RNAs (0.5 µg) were reverse transcribed using iScript Reverse Transcription Supermix (Biorad) containing randoms hexamers. The resulting cDNAs (2.5ng) were mixed with 5x SsoAdvanced Universal SYBR Green Supermix (Biorad) and gene-specific primer pairs obtained from Qiagen. Real-time qPCR reactions were then run on a CFX96 Real-Time System C1000 Thermal Cycler (Bio-Rad, Marnes-la-Coquette, France). All samples were analyzed in triplicate and normalized for B2m and PGK1 expression. The reaction mixture was subjected to real time PCR cycling starting with hot start followed by 40 denaturing cycles at 95 °C for 3 s and annealing extension at 60 °C for 30 s. A final dissociation step was included to determine uniformity of product melting temperature. Data analysis was performed with the CFX Manager Software, 2.1 (Bio-Rad, Marnes-la-Coquette, France), which incorporates the variability of data from both the housekeeping and target genes (*pdgfrα, mbp, Pak1*) to calculate statistical significance. All samples were assayed in triplicate for each target or reference gene and the averaged values were used as Cycle Threshold (Cq). Changes in the relative expression of genes of interest (ΔCq) were calculated according to a normalization to the endogenous controls B2m and PGK1 and then the ΔΔCq was calculated.

### Western Blot

Cultured cells were lysed at 4°C in RIPA buffer (150 mM NaCl, 1.0% IGEPAL^®^ CA-630, 0.5% sodium deoxycholate, 0.1% SDS, 50 mM Tris, pH 8.0, Sigma-Aldrich, St. Quentin Fallavier, France) with the addition of 1 mM DL-Dithiothreitol (Sigma-Aldrich, St. Quentin Fallavier, France), and a protease/phosphatase inhibitor cocktail (Sigma-Aldrich, St. Quentin Fallavier, France). Protein extracts were denatured 5 min at 95°C in a Laemmli buffer (Bio-Rad, Marnes-la-Coquette, France) with β-mercaptoethanol (1/10, Sigma-Aldrich, St. Quentin Fallavier, France). Protein concentration was determined with the Micro kit BCA Protein Assay (Thermo Scientific, Courtaboeuf, France, Courtaboeuf, France). A total of 20 μg of proteins were analyzed in a polyacrylamide gel (4–20% Mini-PROTEAN TGX Precast Protein Gels, Bio-Rad, Marnes-la-Coquette, France) and separated by electrophoresis. Then proteins were transferred into a PVDF membrane (Bio-Rad, Marnes-la-Coquette, France) with a transfer buffer (Tris 20 mM, Glycine 150 mM, Ethanol 20%). The membranes were incubated for 1 hr in a saturation solution (TBS [Tris 20 mM, Sodium Chloride 150 mM, pH 7.6] + Tween20 0.1% + dry milk 5% or 5% BSA) then incubated with primary antibodies overnight at 4°C and washed three times in TBST (TBS+ 0.1% Tween20). The primary antibodies were revealed using secondary antibodies coupled to HRP (horseradish peroxidase (45min at RT)). HRP is detected by a chemiluminescence assay using the kit Clarity Western ECL Substrate (Bio-Rad, Marnes-la-Coquette, France), whose light emission is captured by the camera of the device ChemiDoc Touch (Bio-Rad, Marnes-la-Coquette, France). Quantification of relative protein expression was performed with Image Lab 6.0 software (Bio-Rad, Marnes-la-Coquette, France).

Primary and secondary antibodies used for western blots and immunostaining are listed in **Sup. Table 3**.

### Immunoprecipitation

Primary rat OPCs were lysed at 4 and 5DIV of differentiation as described above. Total amount of proteins was quantified and 250µg of cell lysates were immunoprecipitated overnight with 0,1µg of anti-PAK1 or 0,7µg of anti-Merlin (see Sup.Table 3) at 4°C on a circular shaker, 20 rpm. Incubation with pre-cleaned A/G protein magnetic beads (ThermoFischer Scientific, Courtaboeuf, France) was then initiated for 2h, 4°C, on a circular shaker, 20 rpm. Samples were resolved by electrophoresis and Western blotting as previously described after several washing and denaturation steps.

### Actin ratio

To measure F- and G-actin amounts in cells, we used the G-actin/F-actin *in vivo* assay kit (Cytoskeleton Inc., Denver, USA) according to manufacturer’s protocol. A duplicate of 500μg of proteins were collected for each experimental condition prior to the ultracentrifugation. Equal volumes of the purified fractions were analyzed on a polyacrylamide gel (4–20% TGX Precast Gels, Bio-Rad, Marnes-la-Coquette, France). Total actin was then determined by immunoblotting actin for each experimental condition. F-actin was then rationalized on total actin (F+G-actin).

### Cerebellar slice culture

Cerebellar organotypic cultures were prepared from newborn (P0) mice as previously described previously (60). Brains were dissected in cold Gey’s balanced salt solution (Invitrogen, Villebon-sur-Yvette, France) supplemented with 5 mg/mL glucose. Cerebellar parasagittal slices (350 μm thick) were cut on a McIlwain tissue chopper and transferred onto 30 mm diameter Millicell culture inserts with 0.4 mm pores (Merck, Fontenay sous Bois, France). Slices were maintained in culture in six-well-plates containing 1 mL of nutrient medium per well, at 37°C, under a humidified atmosphere containing 5% CO2. The serum medium consisted of 50% basal medium with Earle’s salts (BME, Invitrogen, Villebon-sur-Yvette, France), 25% Hanks’ balanced salt solution (Invitrogen, Villebon-sur-Yvette, France), 25% horse serum (Invitrogen, Villebon-sur-Yvette, France), l-glutamine (1 mM), 5 mg/mL glucose, 0.2% of antibiotics (Penicillin–Streptomycin [10,000 U/mL], Sigma-Aldrich, St. Quentin Fallavier, France).

### NVS-PAK1-1 treatment

Optimal concentration of NVS-PAK1-1 (Tocris Bioscience, Noyal Châtillon sur Seiche, France) was determined by applying serial dilutions (125nM to 1µM) on rat OPCs from 0DIV to 4DIV of differentiation. A concentration of 250nM was selected to efficiently inhibit PAK1 kinase activity, compared to control condition (Dimethylsulfoxyde; DMSO). This concentration was applied from 0 to 3DIV or from 0 to 6DIV of differentiation. The same concentration of NVS-PAK1-1 was applied on P0 cerebellar slices from 5 to 10DIV.

### siRNA knockdown experiments

A combination of 4 siRNAs targeting *Nf2* as well as a combination of 4 control siRNAs was used for this experiment (Accell siRNA, Horizon Discovery, Cambridge, UK). Optimal concentration of si*Nf2* was determined by applying serial dilutions (100nM to 1 µM) on rat OPCs from 0 to 4DIV of differentiation. A concentration of 200nM was selected to efficiently downregulate Merlin expression, compared to control condition. This concentration was applied to rat OPCs during the first 3DIV of differentiation, without changing the medium. After this period, the medium was changed, and protein extraction or immunostaining was initiated at 5DIV. For the NVS rescue experiment, rat OPCs were subjected to the same protocol, with the difference that from 3DIV, treatment with NVS-PAK1-1 (250nM) or DMSO was applied between 3 and 6DIV of differentiation.

### Cerebellar slices immunostaining

Cerebellar slices were fixed with 2% of paraformaldehyde (PFA, Electron Microscopy Sciences Nanterre, France) for 1 hr at room temperature (RT) and washed several times with PBS. Slices were immersed in Clark’s solution (95% ethanol/5% acetic acid) for 20 min at 4°C and were then washed several times with PBS. All slices were incubated during 1 hr in PBS containing 4% bovine serum albumin (BSA)/4% donkey serum/0.2% triton-X100. Then, they were incubated with the primary and secondary antibody in PBS/2% BSA/2% normal donkey /0.1% triton X-100 solution overnight at 4°C and 2 hr at RT, respectively. Three washes with PBS/0.1% Triton X-100 were performed after each incubation. Last, slices were mounted in Fluoromount-G (SouthernBiotech, Birmingham, USA).

### Oligodendrocyte culture immunostaining

OLs were fixed with 2% of PFA in 1X PBS for 10 min at RT. After several washes of PBS, cells were permeabilized with 0.05% Triton-X100 (15 min at RT) before incubation with primary and secondary antibodies in PBS/1% BSA/5% donkey or goat serum solution, for respectively, 90 and 45 min at RT. Several washes in PBS were performed after incubation with primary and secondary antibodies. Coverslips were mounted in Fluoromount-G (SouthernBiotech, Birmingham, USA).

### Brain section preparation and immunostaining

Animals were deeply anesthetized with pentobarbital (200 mg/kg) and transcardially perfused with 2% PFA in 1X PBS. Fixed brains were dissected and post-fixed overnight in 2% PFA at 4 °C. Samples were immersed in 15% sucrose in 1X PBS for 24 h and subsequently immersed in 30% sucrose in 1X PBS for 48 h. Samples were then embedded in Tissue-Tek O.C.T. (Sakura, Torrance, USA) compound and frozen on liquid nitrogen. Brains were sliced coronally in 12 μm sections on a cryostat, transferred onto glass slides (Superfrost Ultra Plus, Fisher), and stored at −80 °C. Samples were immersed in PBS 1X for 15min and then blocked in 1% BSA/5% Donkey serum /0.2% Triton-X100 for 1 h. Antibodies were diluted in 1% BSA/5% Donkey serum /0.025% Triton-X100. Samples were incubated with primary antibodies overnight at 4°C. They were then washed with PBS 1X/0,025% Triton-X100 under agitation and incubated in the dark with the secondary antibodies for 1hr at RT. Slides were rinsed with PBS 1X/0,025% Triton-X100 under agitation, mounted in Fluoromount-G (SouthernBiotech, Birmingham, USA) with coverslips and left overnight to dry at 4°C.

### Image acquisition

Fluorescence imaging was performed with a 20X objective using Axioscan.z1 microscope slide scanner (Zeiss, Rueil Malmaison, France), enabling *in vitro* and *in vivo* quantification of oligodendroglial differentiation. Image processing and analysis were performed using ZEN 10.0 software (Zeiss, Rueil Malmaison, France). Images of cerebellar slices were acquired with a confocal SP8 X white light laser (Leica, Nanterre, France), at ×40 magnification. Images for PAK1 expression were acquired with the same confocal setting (60x magnification) and STED nanoscope STEDYCON (100x magnification) (Abberior, Göttingen, Germany) for super-resolution images.

### Quantification of differentiation rate *in vitro*

Oligodendrocytes were immunolabelled with Sox10 (OPCs and OLs) and MBP (OLs). The differentiation rate was defined as Sox10+/MBP+ cells on Sox10+ cells and was determined manually for at least 400 cells for DMSO/NVS-PAK1-1, WT versus PAK1 KO and si*Nf2* treatment and rescue conditions. At least 3 experiments were analyzed per treatment.

For lentiviral infection conditions, the differentiation rate was defined as Sox10+/GFP or HA+/MBP+ on Sox10+/GFP or HA+ cells and was determined manually for at least 50 cells per culture.

### Quantification of myelin membrane area

Surface membrane area of each individual oligodendrocyte was measured manually by delimiting the periphery of Sox10+/MBP+ cell’s membrane. Oligodendrocytes in contact with other OLs or with microglia/astrocytes were excluded and only isolated oligodendrocytes were included in the quantification. Quantified surface was measured in at least 3 experiments per treatment. In order to consider surface membrane area of transduced/induced cells, oligodendrocytes treated with lentiviral vectors were immunolabelled with Sox10, MBP and GFP for GFP vector, or HA for PAK1 mutant conditions. Surface membrane of individual OLs was therefore determined on Sox10+ MBP+ GFP+ or Sox10+ MBP+ HA+ cells.

### Quantification of myelination and differentiation index *ex vivo*

Both indexes were calculated with Fiji (ImageJ) macros as previously described in (60). At least 3 independent experiments were analyzed per treatment.

### Quantification of differentiation rate *in vivo*

Positive cells were counted semi-automatically with a homemade macro on ImageJ (Fiji), in at least 3 adjacent coronal sections of at least 3 different animals. A region of interest (ROI) was manually determined and the number of Olig2+, Olig2+/ CC1+ cells was quantified at P60. For *Pak1*cKO mutant sections, oligodendroglial GFP+ cells were considered in the quantification. We selected sections starting from the first rostral section where the corpus callosum has been completely formed.

### Electron microscopy experiments and analysis

Animals were deeply anaesthetized with pentobarbital (200 mg/kg) and transcardially perfused with fresh 4% glutaraldehyde/5% paraformaldehyde in 0.1 M phosphate buffer and post-fixed 1hr in the same solution. Brains were cut in 100 μm coronal slices on the vibratome. Slices containing the regions of interest were post-fixed in 2% osmium tetroxide, dehydrated in graded ethanol and flat embedded in epoxy resin. The flat embedded regions were cut transversely under a stereomicroscope. Sections were cut to two or three different levels of depth in the block. Ultra-thin sections of 70 nm were stained with lead citrate and examined with a Hitachi 7000 electron microscope (EM). Samples were imaged at 44.0 k× for quantifications of axon diameter and g-ratio and at 11.0 k×, for the quantification of the density of myelinated axons. G-ratio was calculated as described in (60). EM pictures were used to estimate the g-factor using MRI g-ratio toolset (“MRI g-ratio Tools ImageJ macro” 2014). Only transverse axons surrounded by a compact myelin were analyzed. For each animal, at least 50 axons were analyzed manually using Fiji (ImageJ) to determine the g-ratio and myelinated axon density.

### Cell sorting of primary OPCs

For proteomics analysis, OPC were purified by magnetic cell sorting (MACS). Brains were collected from wistar rat at P5 and cortices and corpus callosum were dissected and dissociated using neural tissue dissociation kit (P) (Miltenyi Biotec, Paris, France) with gentle MACS Octo Dissociator (Miltenyi Biotec, Paris, France). OPC magnetic cell sorting was performed using anti-O4-coupled-beads (O4 MicroBead Kit, Miltenyi biotec, Paris, France) and MultiMACS Cell24 Separator Plus (Miltenyi biotec, Paris, France). 5.10^5^ cells were plated on petri dish coated with polyethyleneimine (PEI, 100µg/ml, Sigma-Aldrich, St. Quentin Fallavier, France) and amplified in a proliferating medium. At 2DIV, OPCs were placed in differentiation medium until 5DIV. Cells were lysed at 4°C with RIPA buffer and protein concentration was determined with the Micro kit BCA Protein Assay (Thermo Scientific, Courtaboeuf, France, Courtaboeuf, France).

### PAK1 immunoprecipitation and proteomics analysis

Lysates from oligodendrocyte cultures were prepared at 5 DIV using RIPA buffer as described above. We used 500µg of proteins for PAK1 immunoprecipitation and we followed the protocol described above. After several washing and denaturation steps, each immunoprecipitation of PAK1 was validated by western blot prior proteomics analysis. After SDS-PAGE of eluates samples (few millimeters migration) and silver staining of the gel, the small lanes were excised and then cut into small 1mm^3^ pieces. Gel pieces were destained with a solution of 50 mM ammonium bicarbonate (AmBic) and 50% ethanol (EtOH) at 60°C. This step was followed by reduction (incubation in 10 mM DTT in 50 mM AmBic for 30 min at 56°C), and alkylation (incubation in 50 mM iodoacetamide in 50 mM AmBic for 30 min at RT in the dark). Proteins were digested with 200 ng trypsin per well overnight at 37°C (in-gel tryspin digestion). The gel pieces were washed twice in 60% acetonitrile (ACN) / 0.1% trifluoroacetic acid (TFA) for 20 min. Peptide extracts were then dried using a Speed-Vac and resuspended in 20 μL of 2% ACN / 0.1% formic acid (FA). Sample desalting was performed with home-made StageTips consisting in a stack of two reverse-phase C18 layers (Empore SPE Disks C18, Sigma Aldrich) inserted in a 10 μL tip. This step was carried out using the Digest-ProMSi robot (CEM). Each StageTip was first hydrated with methanol, activated with a 50% ACN / 0.5% acetic acid (HAc) solution and then with a 0.5% HAc solution. The peptide solution previously diluted with 0.5% HAc to a final volume of 70 μL was then loaded to the StageTip. The peptides retained in the StageTip were washed using the 0.5% HAc solution. Finally, the peptides were eluted with a 80% ACN / 0.5% HAc solution, completely dried (Speed-Vac) and resolubilized in 20 μL of 2% ACN / 0.1% FA. Samples were stored at –80◦C until mass spectrometry (MS) analysis.

Peptide samples were analyzed with a nanoElute UHPLC (Bruker) coupled to a timsTOF Pro mass spectrometer (Bruker). Peptides were separated on an analytical column RP-C18 Aurora2 (25 cm, 75 μm i.d., 120 Å,1,6 µm particle size, IonOpticks) at a flow rate of 200 nL/min, at 45°C, with mobile phase A (FA 0.1 %) and B (ACN 99.9% / FA 0.1%). The elution gradient was run as follows: from 2% to 5% B in 1 min, from 5% to 13% B in 18 min, from 13% to 19% in 7 min and from 19% to 22% B in 4 min. MS acquisition was run in DDA mode with PASEF. Accumulation time was set to 180 msec in the TIMS tunnel. Capillary voltage was set at 1,6 kV, mass range from 100 to 1700 m/z in MS and MS/MS. The range of ion mobility window was set to 0.7-1.2 1/K0. The quadrupole isolation width was 2 Th below 700 m/z, ramped from 2 to 3 Th between 700 and 800 m/z and remained at 3 Th above 800 m/z. Dynamic exclusion was activated for ions within 0.015 m/z and 0.015 V.s/cm² and released after 0,4 min. Exclusion was reconsidered if precursor ion intensity was 4 times superior. Low abundance precursors below the target value of 14,000 a.u and intensity of 1,000 a.u. were selected several times for PASEF-MS/MS until the target value was reached. Parent ion selection was achieved by using a two-dimensional m/z and 1/K0 selection area filter allowing the exclusion of singly charged ions. The total cycle time was 1,3 sec with 6 PASEF cycles. Raw data were processed using Data Analysis 5.3 (Burker) to generate mgf files.

For protein identification, mgf files were processed with X!Tandem pipeline 0.4.42 (73) using X!Tandem engine 2017.2.1.4. The search parameters were as follows: mass tolerance of 20 ppm for MS1 and MS2, carbamidomethyl as fixed modification and oxidation (on M residue) and protein N-ter acetylation as variable modifications; 1 missed cleavage allowed. Searches (normal and decoy) were performed against the combination of two databases: UniProt rattus norvegicus database (UP000002494_10116.fasta reference proteome, 2021-02 release, 21 594 entries) and a contaminants database. The p-value of peptides and proteins were adjusted to get corresponding FDR < 1%, with a minimum of 2 peptides per protein.

For protein quantification, Bruker raw files were analyzed using MaxQuant software 2.1.3.0 (73) against the UniProtKB *rattus norvegicus* proteome database (UP000002494_10116, 21 594 entries, release 2021_02). The following parameters were used for peptides and proteins identification: trypsin as enzyme specificity with a maximum of two missed cleavages; cysteine carbamidomethylation as a fixed modification and protein N-terminal acetylation and methionine oxidation as variable modifications; peptides minimum length of seven amino acids; precursor mass tolerance of 20 ppm (first search) and 10 ppm (main search); fragment mass tolerance of 40 ppm; 1% FDR. Label-free protein quantification (LFQ) was performed using “unique+razor” peptides with no normalization, no matching between runs (MBR) and with the “min ratio count” parameter set to 1.

The output protein file was processed with ProStar 1.30.7 (74) to keep only proteins detected in at least 3 samples among 5 in at least 1 of the 2 conditions. Missing values were imputed using SLSA (Structured Least Square Adaptative) algorithm for partially missing values in each condition and DetQuantile algorithm for missing values in an entire condition. To select PAK1 relevant binding partners, data were statistically processed using limma test and filtered to retain only differentially expressed proteins (FDR 5%) with a fold change ≥ 3 between IP-PAK1 and IP-IgG conditions.

Gene Set Enrichment Analysis (GSEA) was performed by ClusterProfiler package (75) in RStudio 2023.06.0 and R 4.3.1 softwares. Proteins list was submitted to ClusterProfiler using the following parameters: org.rn.eg.db 3.17.0 database, p-value adjusted cutoff = 0.05 using the Benjamini-Hochberg (BH) method, q-value cutoff = 0.05, minimal size of genes = 3, maximal size of genes = 500. The dot plot method was used to visualize the enriched Gene Ontology (GO) terms.

### Illustrations and statistics

Figure panels were prepared using Fiji (ImageJ) and Adobe Photoshop version 10.0 software (Adobe System, Inc). Statistical analyses were performed with Rstudio (version 2023.06.1+524) (https://cran.r-project.org/) using “ggplot2” package to display results as boxplots, violin plot and scatter plot. Shapiro– Wilk or Kolmogorov-Smirnov tests were used to check for data normality. If failed, when comparing more than two groups of variables, unpaired Kruskal–Wallis test was performed followed by a by Dunn post-hoc test and p-value displayed, while unpaired bilateral Wilcoxon–Mann–Whitney test was performed between each pair of variables and p-value only displayed when significant. Variables that passed the normality test were analyzed by unpaired Student’s t test for comparing two groups. The statistical tests and the number of experiments are described in each figure legend. Some of the statistical analyses were also performed with MatLab (https://www.mathworks.com) to display results as histogram and matrix correlation for western blot analysis. Permutation tests (M=1000) were done to analyze differences between western blot conditions and Pearson correlation coefficients were established between each protein of interest. We further investigated pairwise protein relations when relevant and fitted linear models to the data.

## Acknowledgements

We are grateful to Dr. Hyroyushi Miyoshi (Keio University, Japan) and the Riken BioResources for providing the pCSIV-IRES-hKO vector. All animal work was conducted at the ICM PHENOPARC Core Facility. We thank Nadège Sarrazin and all the members of the PHENO-ICMice for their help. We thank the following core facilities from the Paris Brain Institute-ICM: Histomics, Celis, iVector and ICM-Quant. We would like to thank Asha Baskaran, Dominique Langui for their help with the electron microscopy experiments. We would also like to thank David Akbar from ICM-Quant and Solenne Chardonnier from P3S for their advice. This work was supported by Fondation pour l’Aide à la Recherche sur la Sclérose en Plaques (ARSEP grants n°R18103DD, n°R19233DD, n°R20219DD), Fondation Jérôme Lejeune (n° S.1400.FONDLEJEUNE.1), Fondation Marie-Ange Bouvet-Labruyère (S.1400.NEURATRIS-LB) and Carnot Maturation to L.B-O. This work was also supported by the Paris Brain Institute-ICM and NeurATRIS-joint project to L.B-O and JV.B. L.B. and N.A. are supported by fellowships from the French Ministry of Research (ED158-ED3C). They are also supported by ARSEP and FRM (Fondation pour la Recherche Médicale), respectively. L.B-O is supported by Sorbonne Université. Two figures were created with BioRender.com.

## Author contributions

L.B, N.A and L.B-O conceived and designed the experiments. L.B and N.A carried out the experiments, analyzed the data, prepared the figures, and contributed to the drafting of the manuscript. B.NO and JV.B gave advice and edited the manuscript. L.B, R.BM and Y.V carried out the biostatistical analyses. C.B participated in EM experiments and data analysis. S.G performed qRT-PCR experiments. JV.B and K.D prepared the vectors used in this study. S.H and C.P performed proteomics experiments and analysis. K.K, C.D, R.P and H.H contributed to some experiments. L. B-O supervised the project, wrote, and revised the manuscript.

