## Supplementary material for "PAK1 and NF2/Merlin jointly drive myelination by remodeling actin cytoskeleton in oligodendrocytes": SI Figures & table 3

### Supplementary figures and legends

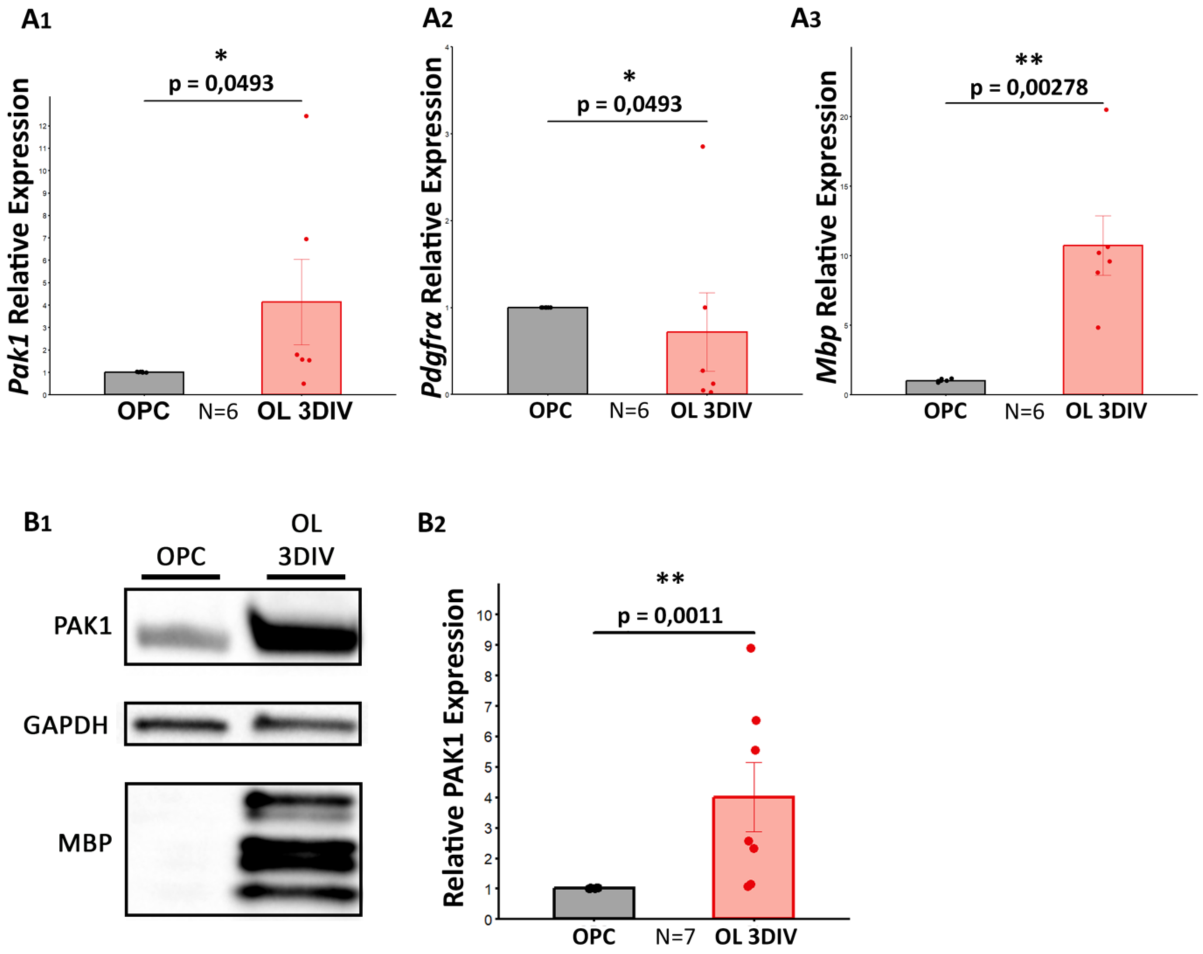


**Supplementary Figure 1. PAK1 expression is increased during OPC differentiation into OLs. (A)** Quantification by RT-qPCR of the relative mRNA level of *Pak1* **(A1)**, *Pdgfra* **(A2)** and *Mbp* **(A3)** between mouse OPCs and OLs at 3DIV of differentiation. Note the significant increase of *Pak1* and *Mbp* in OLs compared to OPCs, and the reduction of *Pdgfrα* in the OLs compared to OPCs. (N= number of replicates, Wilcoxon rank-sum test). **(B1)** Western blot illustration of PAK1 and MBP expression OPCs and OLs at 3DIV GAPDH is used as a loading control. Note the absence of MBP expression in OPCs extracts. **(B2)** Western blot quantification of PAK1 expression rationalized on GAPDH and normalized on OPC condition. Note the significant increase of PAK1 expression in OLs compared to OPCs (N= number of replicates, Wilcoxon rank-sum test).

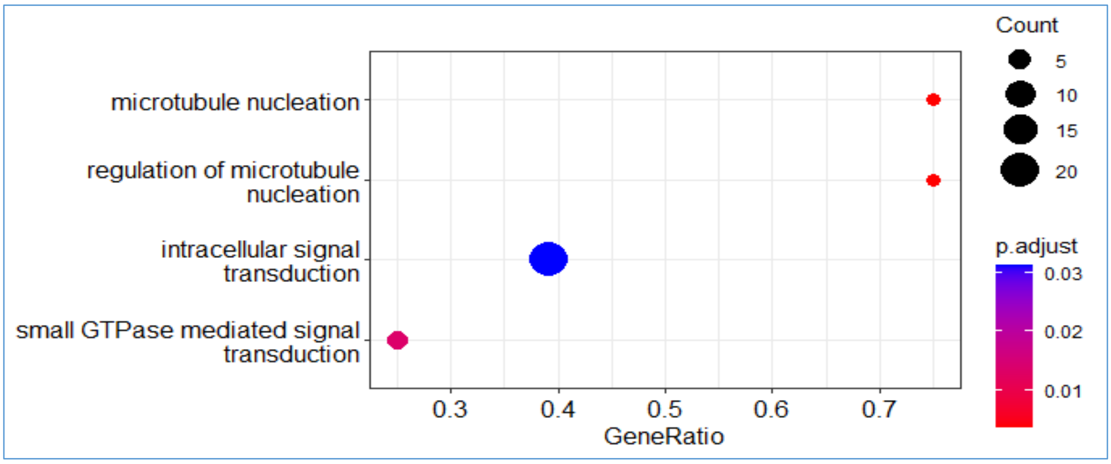


**Supplementary Figure 2.** Dot plot of gene set enrichment analysis (GSEA) performed by ClusterProfiler package. Gene Ontology (GO) terms significantly enriched (FDR < 0.05, p< 0.05) among all identified potential PAK1 binding partners are grouped by the main category-Biological Process (GO:BP). Counts indicate the number of proteins annotated with each term/pathway and dots are colored by ascending normalized enrichment score.

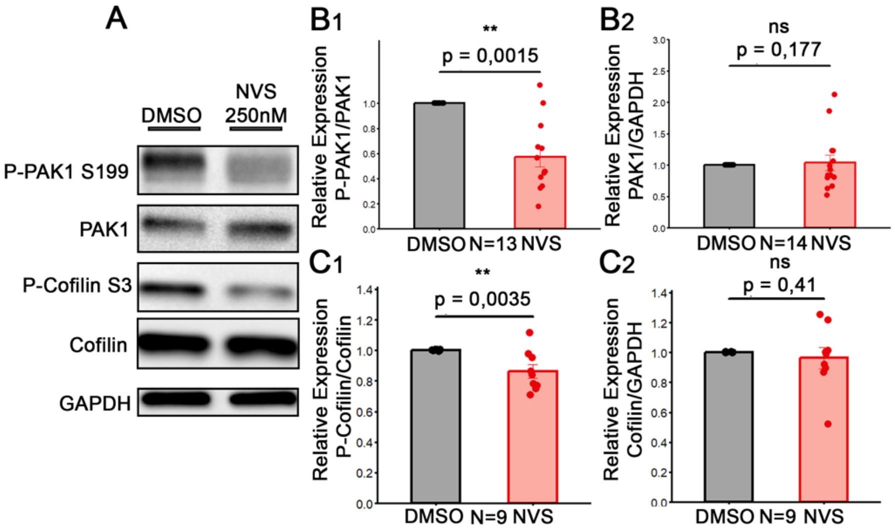


**Supplementary Figure 3. NVS-PAK1-1 treatment efficiently inhibits PAK1 catalytic activity in OL cultures. (A)** Western blot illustration of P-PAK1 (phosphorylated PAK1 on serine 199, S199), PAK1, P-Cofilin (phosphorylated Cofilin on serine 3, S3) and Cofilin expression in OLs treated with DMSO or NVS 250nM and analyzed at 4DIV. GAPDH is used as a loading control. **(B)** Quantification of phosphorylated PAK1 on S199 **(B1)** and PAK1 **(B2)** rationalized on GAPDH and normalized on the control condition. Note the significant decrease of P-PAK1/PAK1 ratio in the NVS condition compared to control, without affecting PAK1 expression (N= number of replicates, Wilcoxon rank-sum test). **(C)** Quantification of P-Cofilin **(C1)** and Cofilin **(C2)** rationalized on GAPDH and normalized on the control condition. Note the significant decrease of P-Cofilin/Cofilin ratio in the NVS condition compared to control, without affecting Cofilin expression (N= number of replicates, Wilcoxon rank-sum test).

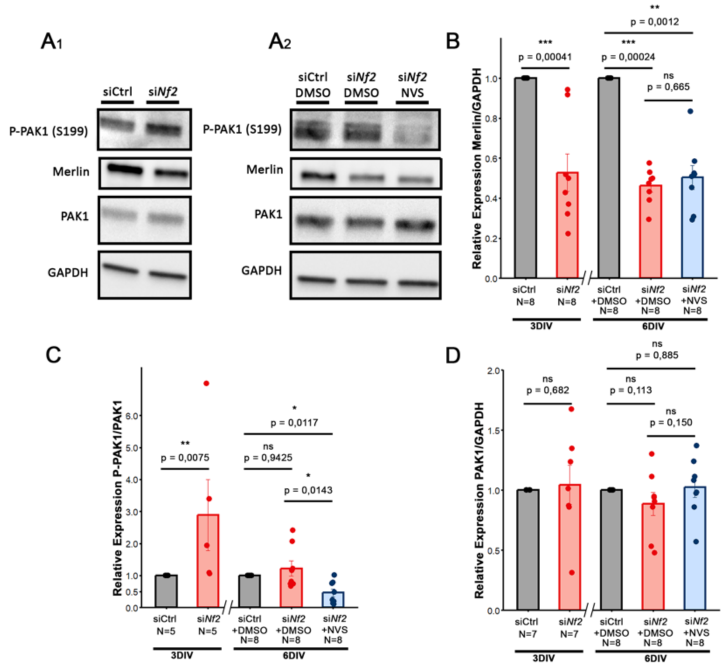


**Supplementary Figure 4. NVS-PAK1-1 reverses PAK1 activation in si*Nf2*-treated OLs. (A1)** Western blot of P-PAK1 (S199), PAK1 and Merlin in 3DIV OLs treated with siControl (siCtrl) or si*Nf2* to assess Merlin expression and the effects on P-PAK1/PAK1 ratio at 3DIV. GAPDH is used as a loading control. **(A2)** Western blot of P-PAK1 (S199), PAK1 and Merlin in 6DIV OLs treated with siControl or siNf2 for 3DIV, and with DMSO or NVS for a further 3 DIV. GAPDH is used as a loading control. **(B)** Quantification of Merlin expression rationalized on GAPDH expression after si*Nf2* treatment. Quantifications are normalized on the control condition at 3 and 6DIV. Note the significant decrease of Merlin in both conditions. At 6DIV, the addition of NVS did not change the expression of Merlin (blue) (N= number of replicates, for experiments at 3DIV: Wilcoxon rank-sum test, for experiments at 6DIV: Kruskal-Wallis followed by Dunn post-hoc test). **(C)** Quantification of phosphorylated PAK1 (S199) rationalized over PAK1 expression. Quantifications are normalized on the control condition at 3 and 6DIV. Note the significant increase of the P-PAK1/PAK1 ratio after 3DIV of si*Nf2* treatment compared to the 3DIV control. Similarly, note the significant decrease of the P-PAK1/PAK1 ration at 6DIV after addition of NVS-PAK1 treatment on si*Nf2* treated cultures (N= number of replicates, for experiments at 3DIV: Wilcoxon rank-sum test, for experiments at 6DIV: Kruskal-Wallis followed by Dunn post-hoc test). **(D)** Quantification of PAK1 expression rationalized over GAPDH expression. Quantifications are normalized on the control condition at 3 and 6DIV. Note the absence of significant differences between conditions (for experiments at 3DIV: Wilcoxon rank-sum test, for experiments at 6DIV: Kruskal-Wallis followed by Dunn post-hoc test).

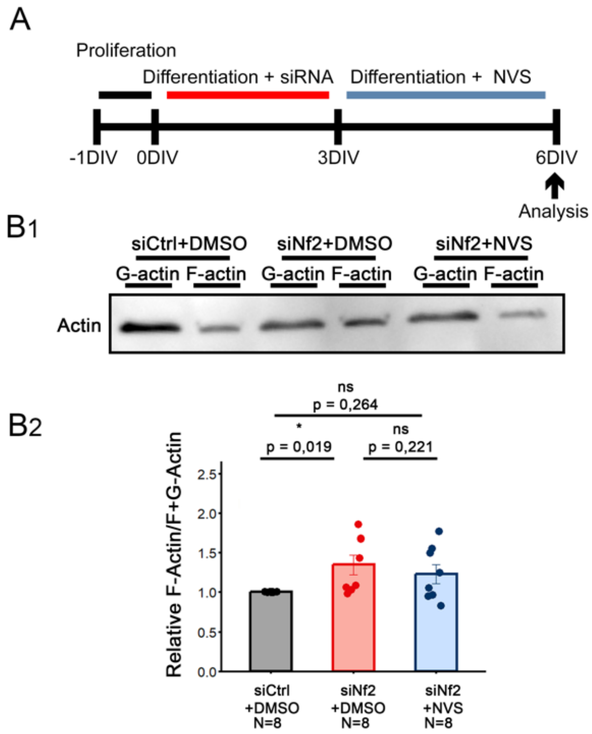


**Supplementary Figure 5. Effects of *Nf2* knockdown in OLs on F and G-actin fractions. (A)** Schematic diagram showing the different experimental steps. After 1 DIV in proliferation medium, OPCs are treated with siRNA (siNf2 or control siRNA (siCtrl)) for 3 DIV in differentiation medium. At 3DIV, medium is changed and DMSO or NVS (250nM) is added. After 3DIV of NVS or DMSO treatment (6DIV of differentiation), cells were analyzed. **(B) (B1)** Western blot analysis of F- and G-actin fractions purified from OLs treated with siRNAs (control or *Nf2*) followed by DMSO or NVS treatment. **(B2)** Quantification of the ratio of F-actin over total actin (F+G-actin) in OLs at 6DIV. It should be noted that, although *Nf2* knockdown leads to an increase in F-actin and NVS treatment corrects this effect (western blot), no statistical differences were observed in F-actin/total actin ratio between the siCtrl and DMSO, si*Nf2* and DMSO, si*Nf2* and NVS conditions (N= number of replicates, Kruskal-Wallis followed by Dunn post-hoc test).

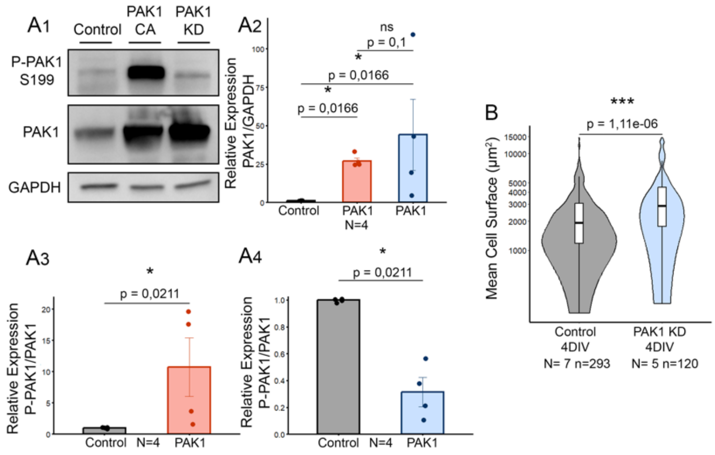


**Supplementary Figure 6. Validation of PAK1 CA et KD vectors, and effects of PAK1 KD on myelin membrane formation. (A) (A1)** Western blot illustration of P-PAK1 (phosphorylated on serine 199, S199) and PAK1 expression in control, PAK1 CA and PAK1 KD transduced HEK 293-T cells. GAPDH is used as a loading control. **(A2)** Quantification of the relative expression of PAK1 in PAK1 CA et KD conditions rationalized on GAPDH and normalized on the control. Note the significant overexpression of PAK1 in both conditions (N=number of replicates, Kruskal-Wallis followed by Dunn post-hoc test). **(A3-A4)** Quantification of relative expression of P-PAK1 in PAK1 CA **(A3)** and PAK1 KD **(A4)** rationalized on PAK1 and normalized on the control condition. Note the significant increase of the P-PAK1/PAK1 ratio in PAK1 CA condition, and its significant decrease in PAK1 KD condition (N=number of replicates, Wilcoxon rank-sum test). **(B)** Cell surface quantification of rat OLs transduced with control or PAK1 KD vectors and analyzed at 4 DIV **(B)**. Note that PAK1 KD overexpression in OLs significantly increases MBP+ membrane surface area at 4DIV compared to controls (N= number of replicates, n= number of cells, Wilcoxon rank-sum test).

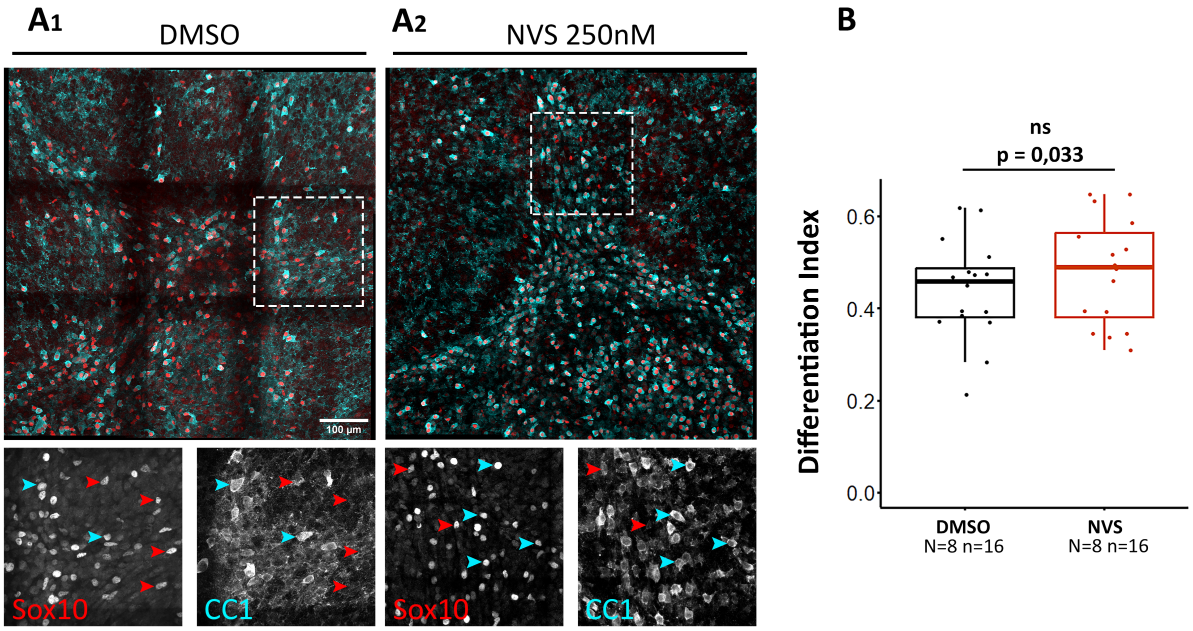


**Supplementary Figure 7. Pharmacological inhibition of PAK1 does not affect OPC differentiation *ex vivo*. (A)** Confocal images of Sox10 (red, oligodendroglial cells) and CC1 (cyan, OLs) immunostainings illustrating OPCs (Sox10+CC1-, red head arrows) and OLs (Sox10+CC1+, blue head arrows) in cerebellar cultures treated with DMSO **(A1)** or NVS 250nM **(A2) at 10DIV**. Two inserts on the bottom of each image represent a zoom in of Sox10 and CC1 straining. **(B)** Boxplot of the quantification of OPC differentiation by calculating the differentiation index. Note the absence of statistical differences in this index for slices treated with NVS compared to controls (N= number of replicates, n= number of cerebellar slices, unpaired Student's t-test).

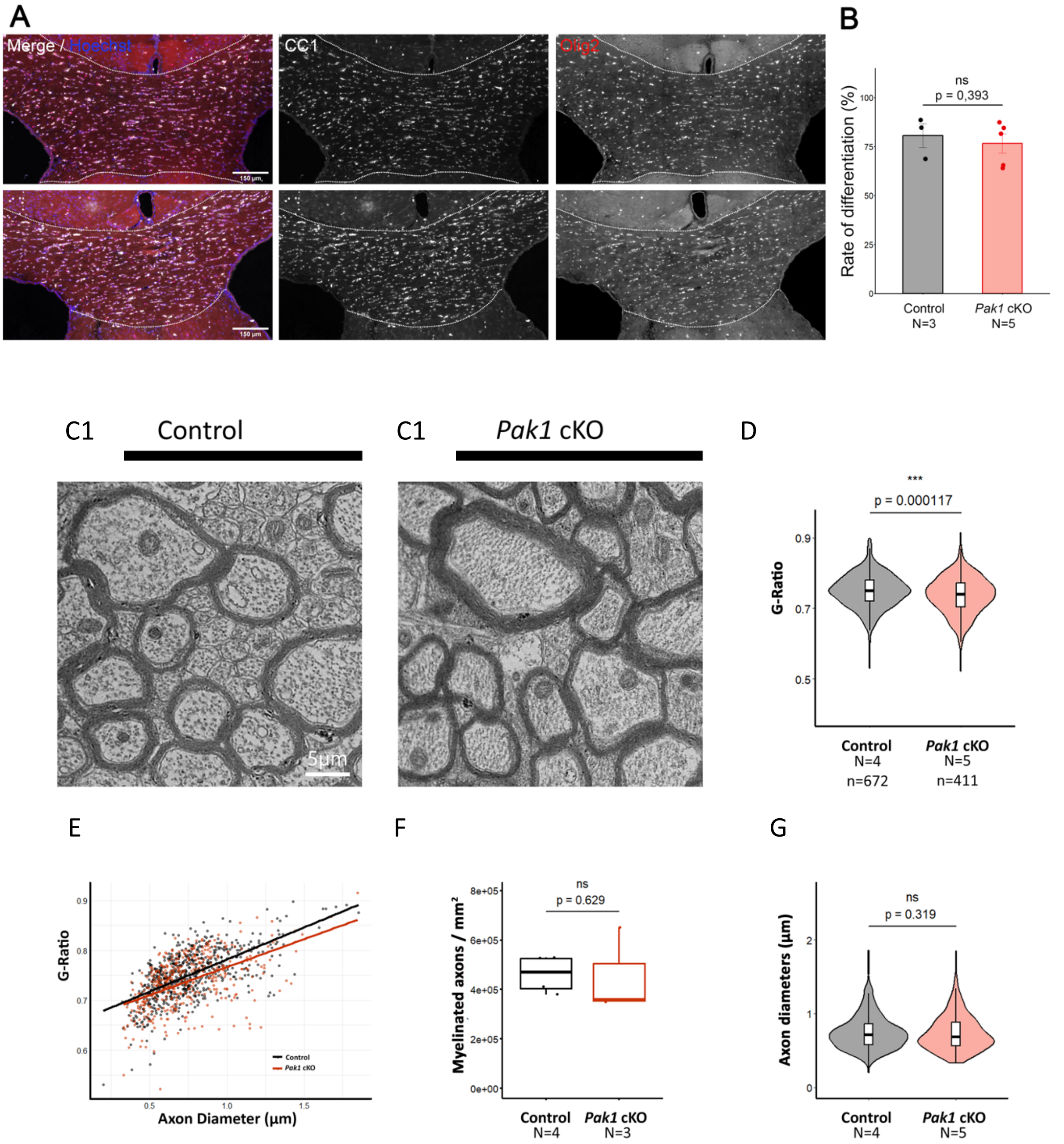


**Supplementary Figure 8. Myelin thickness is increased in the corpus callosum of *Pak1* cKO adult brain.** **(A)** Axioscan imaging of the corpus callosum at P60 in control and *Pak1*cKO animals. The corpus callosum is indicated by the dotted white line. Differentiated OLs (CC1+) are shown in white and total oligodendroglial cells (Olig2+) in red. **(B)** Quantification of the percentage of differentiated OLs (CC1+Olig2+ cells over the Olig2+ cell population) in the corpus callosum at P60 in control and *Pak1*cKO animals. Note that the percentage of differentiated OLs is not different between the two genotypes (Mann-Whitney test). **(C)** Electron micrographs of transverse myelinated axons in the corpus callosum of young adults (P60) control **(C1)** and *Pak1*cKO mice **(C2)**. **(D)** Boxplot of g-ratio analysis in the corpus callosum of control and *Pak1* cKO mice. Note the significant decrease in the g-ratio in *Pak1*cKO animals compared to controls, indicating an increase in myelin thickness in the *Pak1*cKO corpus callosum (N= number of animals, n= number of myelinated axons, unpaired Student's t-test). (**E)** Scatterplots of individual axon diameter measurements as a function of axon g-ratio (each individual data point represents one axon) and linkage to the regression line for each genotype. (**F)** Boxplot of myelinated axon densities. Note that *Pak1* deletion in OLs did not affect the density of myelinated axons (N= number of animals, Wilcoxon rank-sum test). **(G)** Boxplot of axon diameters. Note that *Pak1* deletion in OLs did not affect axonal diameter of neurons (N= number of animals, Wilcoxon rank-sum test).

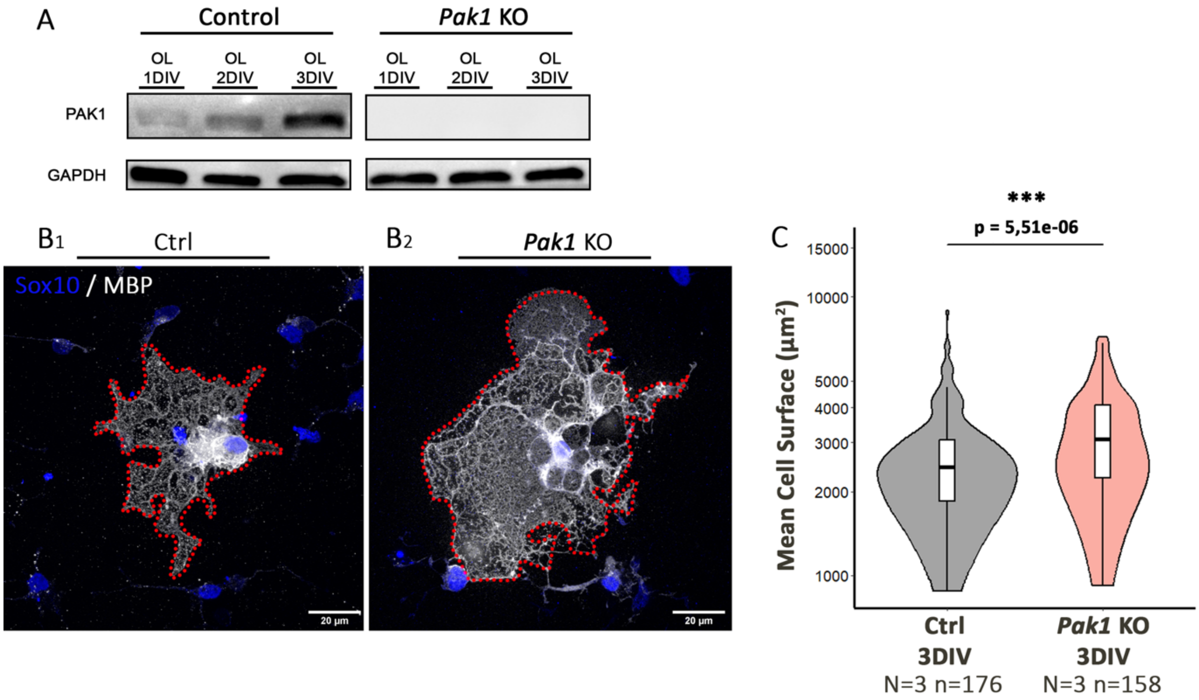


**Supplementary Figure 9. Deletion of *Pak1* in cultured OL promotes myelin membrane expansion. (A)** Western blot illustration of PAK1 expression in mouse OLs (control and *Pak1*KO) at 1, 2 and 3 DIV of culture. In the control OL cultures, PAK1 expression increases with OL maturation. In the *Pak1* KO condition, we validated the absence of PAK1 expression. GAPDH is used as a loading control. **(B)** Confocal acquisition of control and *Pak1*KO mature OLs immunostained with Sox10 (blue, OL lineage), and MBP (white, mature OLs with myelin membrane) antibodies. MBP staining allows the visualization of the shape of the myelin membrane delimited with the red dotted line. **(C)** MBP+ cell surface quantification of control and *Pak1*KO OL cultures at 3DIV. Note the significant increase of mean cell surface in the *Pak1*KO condition compared to the control (N=Number of replicates, n=number of cells, Wilcoxon rank-sum test).

**Supplementary Table.1. A.** List of identified proteins by Xtandem pipline software.
**B.** List of Merlin peptides identified in PAK1 IP.

**Supplementary Table.2.** List of proteins quantified by MaxQuant with 71 statistically significant potential PAK1 binding partners.

**Supplementary Table.3.** List of primary and secondary antibodies.

| **Primary antibodies** | | | | | |
|---|---|---|---|---|---|
| **Target** | **Host** | **Supplier** | **Reference - RRID** | **Dilution** | **Application** |
| MBP | Rat | Abcam | Cat# ab7349; RRID:AB_305869 | 1:200 | IHC/ICC |
| | Rabbit | Cell Signaling Technology | Cat# 78896 ; RRID :AB_2799920 | 1:1000 | WB |
| Calbindin | Rabbit | Swant | Cat# CB38, RRID:AB_10000340 | 1:10 000 | IHC |
| Sox10 | Goat | R&D System | Cat# AF2864, RRID:AB_442208 | 1:50 | IHC/ICC |
| Olig2 | Rabbit | Sigma-Aldrich | Cat# AB9610, RRID:AB_570666 | 1/200 | IHC |
| APC (CC1) | Mouse IgG2b | Millipore | Cat# OP80, RRID:AB_2057371 | 1:400 | IHC |
| MOG | Mouse IgG | Hybridoma | Homemade | 1/10 | WB |
| GFP | Chiken | Aves | Cat# GFP-1020, RRID:AB_10000240 | 1:400 | IHC/ICC |
| Actin | Mouse IgG2b | Cytoskeleton | Cat# AAN02-S | 1/1000 | WB |
| Kusabira-Orange | Rabbit | MBL International | Cat# PM051M, RRID:AB_2876863 | 1:500 | IHC |
| Ha-tag | Rabbit | Cell Signaling Technology | Cat#3724, RRID:AB_1549585 | 1/1000 | ICC |
| P-PAK1 (S199) | Rabbit | Cell Signaling Technology | Cat# 2605, RRID:AB_2160222 | 1/1000 | WB |
| PAK1 | Rabbit | Cell Signaling Technology | Cat# 2602, RRID :AB_330222 | 1/100 | IHC/ICC |
| | | | | 1/1000 | WB |
| CNPase | Mouse IgG1 | Sigma-Aldrich | Cat#C5922, RRID:AB_476854 | 1/400 | ICC |
| Merlin | Rabbit | Cell Signaling Technology | Cat#12888, RRID:AB_2650551 | 1/100 | ICC |
| | | | | 1/1000 | WB |

| | | | | | |
|---|---|---|---|---|---|
| GAPDH | Mouse IgG1 | Sigma-Aldrich | Cat#CB1001, RRID:AB_2107426 | 1/7000 | WB |
| Cofilin | Rabbit | Cell Signaling Technology | Cat#5175, RRID:AB_10622000 | 1/1000 | WB |
| P-Cofilin (S3) | Rabbit | Cell Signaling Technology | Cat#3311, RRID:AB_330238 | 1/1000 | WB |

| **Secondary antibodies and Hoechst** | | | | | | |
|---|---|---|---|---|---|---|
| **Target** | **Host** | **Conjugate** | **Supplier** | **Reference** | **Dilution** | **Application** |
| Goat | donkey | Alexa405 | Abcam | Cat# ab175995, RRID:AB_11011587 | 1 :750 | ICC/IHC |
| Rabbit | donkey | Alexa488 | Thermo Fisher Scientific | Cat# A21206, RRID:AB_2535792 | 1 :1000 | ICC/IHC |
| | | Alexa647 | Thermo Fisher Scientific | Cat# A31573, RRID:AB_2536183 | 1 :500 | ICC/IHC |
| | | HRP | Jackson ImmunoResearch | Cat# 711-035-152, RRID:AB_10015282 | 1/40 000 | WB |
| | goat | Alexa555 | Thermo Fisher Scientific | Cat# A21428, RRID:AB_141784 | 1 :1000 | IHC |
| Mouse IgG total | Donkey | Alexa555 | Thermo Fisher Scientific | Cat# A31570, RRID:AB_2536180 | 1 :1000 | ICC/IHC |
| Mouse IgG2b | Donkey | Alexa647 | Thermo Fisher Scientific | Cat# A31571, RRID:AB_162542 | 1 :1000 | IHC |
| Mouse IgG total | Goat | Alexa647 | Thermo Fisher Scientific | Cat# A21242, RRID:AB_1500900 | 1 :1000 | ICC/IHC |
| Mouse IgG total | Rat | HRP | Jackson ImmunoResearch | Cat# 415-035-166, RRID:AB_2340269 | 1/40 000 | WB |
| Chicken | Donkey | Alexa488 | Jackson ImmunoResearch | Cat# 703-546-155, RRID:AB_2340376 | 1 :1000 | ICC/IHC |

| | | | | | | |
|---|---|---|---|---|---|---|
| | Goat | Alexa488 | Thermo Fisher Scientific | Cat# A11039, RRID:AB_2534096 | 1 :1000 | ICC/IHC |
| Rat | Donkey | Alexa488 | Thermo Fisher Scientific | Cat# A21208, RRID:AB_2535794 | 1 :1000 | ICC/IHC |
| | | Alexa647 | Abcam | Cat# ab150155, RRID:AB_2813835 | 1 :1000 | ICC |
| Hoescht 33258 | | | Sigma-Aldrich | Cat# h21491 | 1 :1000 | ICC/IHC |
